# Manganese depletion leads to multisystem changes in the transcriptome of the opportunistic pathogen *Streptococcus sanguinis*

**DOI:** 10.1101/2020.08.06.240218

**Authors:** Tanya Puccio, Karina S. Kunka, Bin Zhu, Ping Xu, Todd Kitten

## Abstract

*Streptococcus sanguinis* is a primary tooth colonizer and is typically considered beneficial due to its antagonistic relationship with the cariogenic pathogen *Streptococcus mutans*. However, *S. sanguinis* can also act as an opportunistic pathogen should it enter the bloodstream and colonize a damaged heart valve, leading to infective endocarditis. Studies have implicated manganese acquisition as an important virulence determinant in streptococcal endocarditis. A knockout mutant lacking the primary manganese import system in *S. sanguinis*, SsaACB, is severely attenuated for virulence in an *in vivo* rabbit model. Manganese is a known cofactor for several important enzymes in *S. sanguinis*, including superoxide dismutase, SodA, and the aerobic ribonucleotide reductase, NrdEF. To determine the effect of manganese depletion on *S. sanguinis*, we performed transcriptomic analysis on a Δ*ssaACB* mutant grown in aerobic fermentor conditions after the addition of the metal chelator EDTA. Despite the broad specificity of EDTA, analysis of cellular metal content revealed a decrease in manganese, but not in other metals, that coincided with a drop in growth rate. Subsequent supplementation with manganese, but not iron, zinc, or magnesium, restored growth in the fermentor post-EDTA. Reduced activity of Mn-dependent SodA and NrdEF likely contributed to the decreased growth rate post-EDTA, but did not appear entirely responsible. With the exception of the Dps-like peroxide resistance gene, *dpr*, manganese depletion did not induce stress response systems. By comparing the transcriptome of Δ*ssaACB* cells pre- and post-EDTA, we determined that manganese deprivation led to altered expression of diverse systems, including ethanolamine utilization, CRISPR/Cas, and a type IV pilus. Manganese depletion also led to an apparent induction of carbon catabolite repression in a glucose-independent manner. The combined results suggest that manganese limitation produces effects in *S. sanguinis* that are diverse and complex, with no single protein or system appearing entirely responsible for the observed growth rate decrease. This study provides further evidence for the importance of this trace element in streptococcal biology. Future studies will focus on determining mechanisms for regulation, as the multitude of changes observed in this study indicate that multiple regulators may respond to manganese levels.

## 2 Introduction

*Streptococcus sanguinis* is a facultative anaerobe that is typically found in much greater abundance at healthy oral sites than in carious lesions or diseased gingiva (Stingu et al., 2008;Belda-Ferre et al., 2012;Griffen et al., 2012;Gross et al., 2012;Giacaman et al., 2015). The *S. sanguinis* genome encodes a variety of adhesins (Xu et al., 2007;Bensing et al., 2018) that allow it to act as one of the primary colonizers of the salivary pellicle (Socransky et al., 1977;Kolenbrander et al., 2010). It has the capacity to produce (Garcia-Mendoza et al., 1993;Kreth et al., 2008) and survive in (Xu et al., 2014) high concentrations of hydrogen peroxide, which allows it to compete against the dental caries pathogen *Streptococcus mutans* (Kreth et al., 2005). These traits, which have evolved to ensure survival in the highly diverse oral cavity, also make *S. sanguinis* an opportunistic pathogen under the right conditions (Das et al., 2009;Turner et al., 2009b;Bensing et al., 2019). Dental procedures (Kinane et al., 2005;Forner et al., 2006;Lockhart et al., 2008), routine oral hygiene practices (Silver et al., 1977;1979;Moreillon and Que, 2004), mastication (Sreenivasan et al., 2017), and poor oral hygiene (Kholy et al., 2015) can all damage the oral mucosa, allowing bacteria to enter the bloodstream. *S. sanguinis* and certain other bacterial species can bind to cardiac vegetations composed of platelets and fibrin that form on damaged heart valves and endocardium (Lee et al., 2001), leading to the disease infective endocarditis (IE) (Moreillon et al., 2002;Widmer et al., 2006). Recent studies estimate that IE affects more than 40,000 people each year in the U.S. and kills 12-40% (Bor et al., 2013;Cahill et al., 2017;Jamil et al., 2019) due to complications such as congestive heart failure and stroke (Bashore et al., 2006). In the U.S., prevention depends upon antibiotic prophylaxis prior to dental procedures for at-risk patients (Wilson et al., 2007). The economic burden, potential for side effects, and questionable efficacy (Dayer and Thornhill, 2018;Thornhill et al., 2018;Quan et al., 2020) of this practice, as well as the increasing prevalence of antibiotic resistance (Dodds, 2017) are all pressing concerns.

Manganese (Mn) is an essential human micronutrient and has been linked to virulence in many human pathogens, including streptococci (Kehres and Maguire, 2003;Papp-Wallace and Maguire, 2006;Eijkelkamp et al., 2015). Previous work from our lab established that SsaB, the lipoprotein component of the ABC transporter SsaACB, is important for Mn transport and essential for virulence in a rabbit model of IE (Crump et al., 2014). In *S. sanguinis*, Mn acts as a co-factor for superoxide dismutase (SodA) (Parker and Blake, 1988;Poyart et al., 1988) and the aerobic class 1b ribonucleotide reductase (NrdEF) (Makhlynets et al., 2014;Rhodes et al., 2014). Loss of SodA activity alone cannot account for the reduction in virulence (Crump et al., 2014). NrdEF activity is essential for virulence (Rhodes et al., 2014), but it is likely that these are not the only two Mn-cofactored enzymes or Mn-dependent pathways in *S. sanguinis*. In a previous microarray analysis of Mn depletion in *S. pneumoniae* (Ogunniyi et al., 2010), it was found that only a handful of genes were differentially expressed in response to either deletion of the gene encoding PsaA, the pneumococcal ortholog of SsaB, or growth in media without supplemental Mn. However, these data alone are insufficient to explain the decreased growth of these mutants in low-Mn media. In this study, we sought to determine the overall effect of Mn depletion on the transcriptome of *S. sanguinis* in an attempt to identify other Mn-dependent pathways. Here we report that while there were some similarities with this previous study, we found a larger number of differentially expressed genes, providing new insights into the role of Mn in streptococci.

## 3 Results

### 3.1 Selection of fermentor growth conditions for Mn depletion

For this study, we were interested in measuring transcriptional changes resulting from Mn depletion in metabolically active cells. We also wanted to examine the cells as they transitioned from Mn replete conditions to Mn insufficiency, a task that would most easily have been accomplished by addition of a strong and selective Mn chelator to growing cells. However, we were aware of no such chelator. We therefore explored the use of the non-specific chelator EDTA in conjunction with a Δ*ssaACB* mutant. We achieved reproducible, large-scale growth in a fermentor using Brain Heart Infusion (BHI) broth. Typical chemostat conditions (Burne and Chen, 1998) could not be identified that supported growth of the SK36 wild-type (WT) strain but not the Δ*ssaACB* mutant, even when aeration was increased (data not shown). However, we found that when the dilution rate was increased to 0.875 vessel volumes per h, addition of 100 μM EDTA dramatically reduced the optical density (OD_840-910_) of the Δ*ssaACB* mutant cultures (Fig. S1A), while not affecting the WT strain (Fig. S1B). The effect of EDTA addition on the OD of the Δ*ssaACB* cultures typically became apparent after 38 min (Fig. S1 inset). The addition of EDTA slowed the growth of Δ*ssaACB* but did not kill the cells entirely because when the media pumps were shut off ~80 min post-EDTA addition, the OD began to increase immediately (data not shown).

To determine if a lack of available Mn caused the EDTA-dependent reduction in the Δ*ssaACB* growth rate, samples of both WT and Δ*ssaACB* were collected at T_−20_, T_10_, T_25_, and T_50_, where 100 μM EDTA was added to the vessel at T_0_ (Fig. S1). Washed cells were analyzed using inductively coupled plasma optical emission spectroscopy (ICP-OES) (Fig. 1). EDTA addition to WT did not significantly alter cellular levels of any of the four metals measured—Mn, Fe, Zn, or Mg (Fig. 1A). Mn was the only metal significantly reduced in the post-EDTA samples as compared to pre-EDTA for Δ*ssaACB* (Fig. 1B). While iron (Fe) levels were low in the Δ*ssaACB* mutant, they did not drop significantly after the addition of EDTA (Fig. 1B). This is consistent with the metal content of the Δ*ssaACB* mutant measured previously under aerobic growth conditions (Murgas et al., 2020). Neither zinc (Zn) nor magnesium (Mg) levels were significantly affected (Fig. 1). Cobalt and copper levels were at or below the limit of detection in both strains (data not shown).

**Figure 1.**
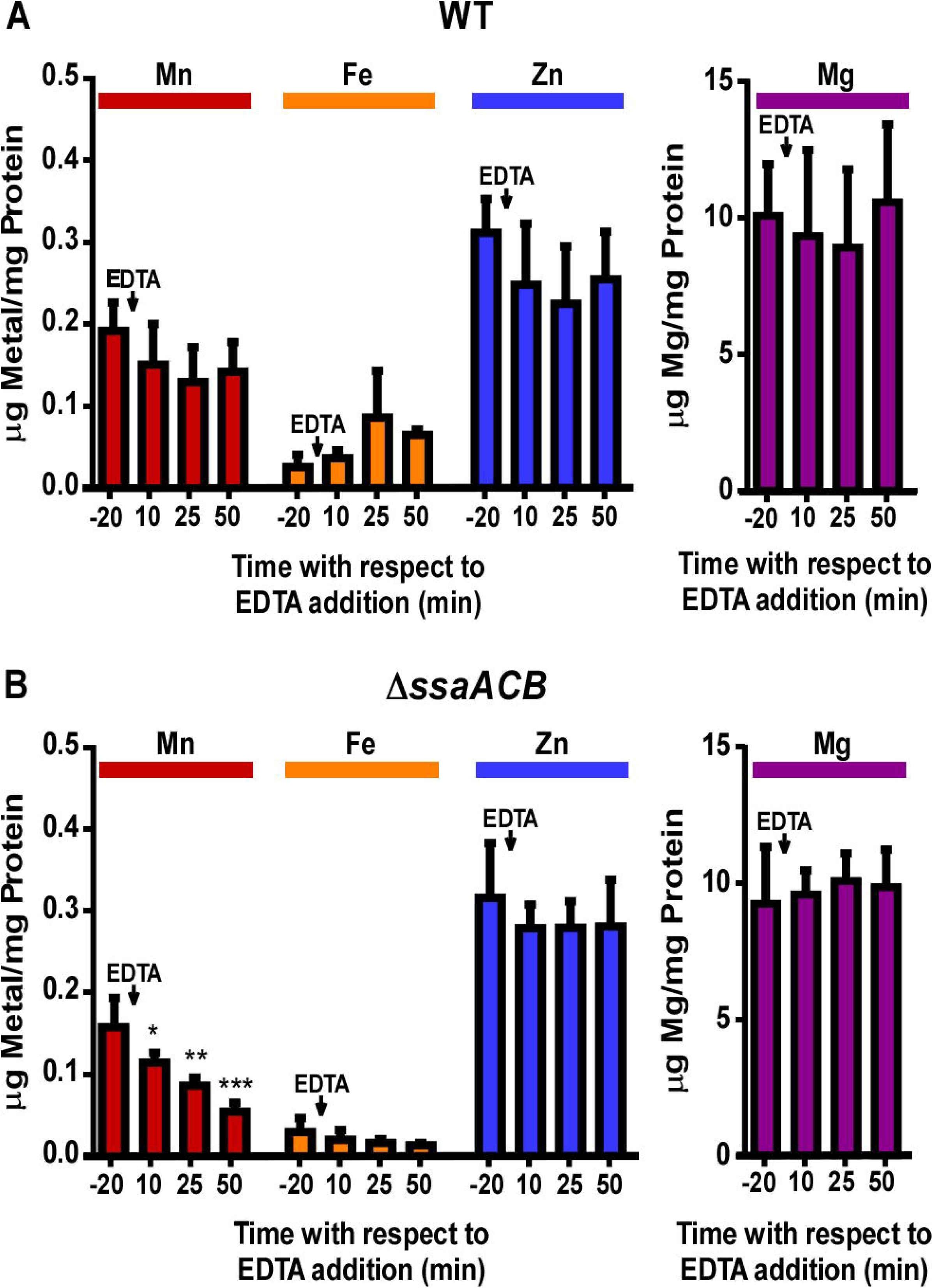
Effect of EDTA on metal content of fermentor-grown WT and Δ*ssaACB* cells. Aerobic fermentor-grown WT (**A**) and Δ*ssaACB* (**B**) cells were collected at each time point and analyzed for cellular metal content using ICP-OES. Means and standard deviations of three replicates are shown. Significance was determined either by repeated measures ANOVA or by one-way ANOVA if matching was not effective, with a Tukey-Kramer multiple comparisons test to T_−20_. \**P* ≤ 0.05, ** *P* ≤ 0.01, *** *P* ≤0.001. Time points not labeled were not significantly different from T_−20_. For Fe, two T_−20_ replicates in (**A**) and at least two replicates for each time point in (**B**) were below the limit of detection.

As another test of metal specificity, 100 μM of either Mn^2+^ or Fe^2+^ sulfate was added to the vessel 70 mins post-EDTA addition. The addition of Mn(II)SO_4_ eliminated, and then reversed, the post-EDTA decline in OD, while Fe(II)SO_4_ had no discernible effect (Fig. S2). The metal content of samples collected 10 mins after addition of Mn(II)SO_4_ or Fe(II)SO_4_ (T_80_) revealed that both Mn and Fe were taken up by cells, resulting in significantly higher levels than at T_−20_ (Fig. S3). Although neither Zn nor Mg levels were significantly affected by addition of EDTA (Fig. 1), 100 μM of either Zn(II)SO_4_ or Mg(II)SO_4_ was added at T_70_ for at least two fermentor runs each and, like Fe(II)SO_4_, neither produced any apparent effect (data not shown).

### 3.2 Overview of transcriptional response of *S. sanguinis* to Mn depletion

In order to assess the impact of Mn depletion on the *S. sanguinis* transcriptome, RNA sequencing (RNA-seq) analysis was performed on Δ*ssaACB* fermentor samples collected at the same time points as above (Fig. S1A). Principal component analysis (PCA) revealed that the samples from each time point clustered together, indicating minimal variation between independent replicates (Fig. 2A). The T_10_ samples overlapped slightly with T_−20_, indicating few early changes in gene expression. The dissimilarities of the RNA-seq profiles were greater at T_25_ and T_50_, suggesting that EDTA treatment increasingly affected the gene expression of Δ*ssaACB* during the tested period.

**Figure 2.**
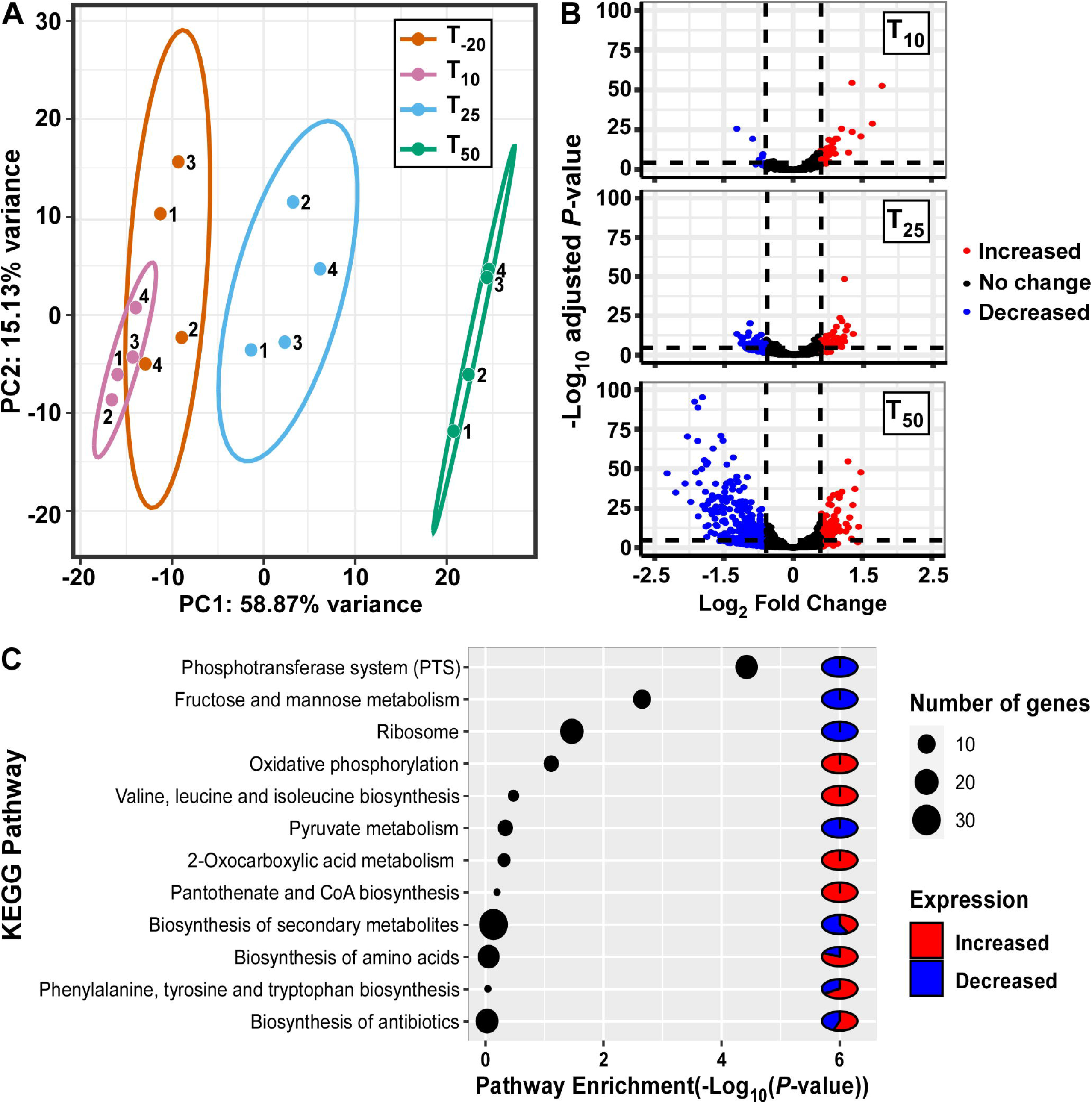
Analysis of gene expression in fermentor-grown *S. sanguinis* Δ*ssaACB* mutant cells. (**A**) Principal component analysis of the RNA-seq samples as determined by the pcaExplorer package for R. Replicates are labeled by fermentor run number. Ellipses are drawn around the 95% confidence interval for each time point. (**B**) Volcano plots comparing each post-EDTA time point to T_−20_ were generated using DESeq2 analysis in the EnhancedVolcano package for R. Genes that are upregulated in the post-EDTA time point are positive on the x-axis (right) and those that are downregulated are negative (left). Genes exhibiting log_2_ fold change > |1| are depicted by either red (> 1) or blue (< 1) spheres. (**C**) Pathway enrichment analysis of differentially expressed genes at T_50_ using DAVID classification of genes and KEGG annotations.

Volcano plot analysis of differentially expressed genes (DEGs; defined as |log_2_| ≥ 1, adjusted *P*-value ≤ 0.05) comparing post-EDTA time points to the pre-EDTA time point revealed that there were only 48 (2.1%) and 139 (6.1%) DEGs at T_10_ and T_25_, respectively (Fig. 2B). In contrast, at 50 mins post-EDTA, 407 genes (17.9%) were differentially expressed, with a number of genes more severely down-regulated (Fig. 2B). Consistent with these results, the growth rate of Δ*ssaACB* decreased dramatically between T_25_ and T_50_ (Fig. S1A).

Gene classification analysis of DEGs at T_50_ using KEGG annotations revealed that involved in sugar metabolism were highly enriched, with phosphotransferase systems (PTS), fructose/mannose transport, and pyruvate metabolism all in the top 12 most highly enriched pathways (Fig. 2C). Other highly enriched pathways included amino acid metabolism, ribosomes, oxidative phosphorylation (ATP synthases), and biosynthesis of coenzymes and various secondary metabolites (Fig. 2C).

RNA-seq trends for several genes of interest with moderate to high expression level changes were validated by measuring mRNA levels of fermentor samples via quantitative reverse transcriptase polymerase chain reaction (qRT-PCR) (Fig. S4). The relative expression levels observed in the qRT-PCR experiments largely replicated the trends observed in the RNA-seq analysis.

### 3.3 Regulation of metal transport genes

As seen in Figure 1, Mn was the only tested metal whose cellular concentration was decreased upon addition of EDTA to Δ*ssaACB* cells. To further investigate the impact of EDTA on metal transport, we examined the expression of metal transport genes (Fig. 3). The kanamycin (Kan) resistance gene *aphA*-3 that replaced the Mn transporter operon, *ssaACB*, in this mutant strain was upregulated in all three post-EDTA time points (Fig. 3). This is consistent with previous results from our lab showing Mn-dependent repression of SsaB expression as measured by western blot (Crump et al., 2014).

**Figure 3.**
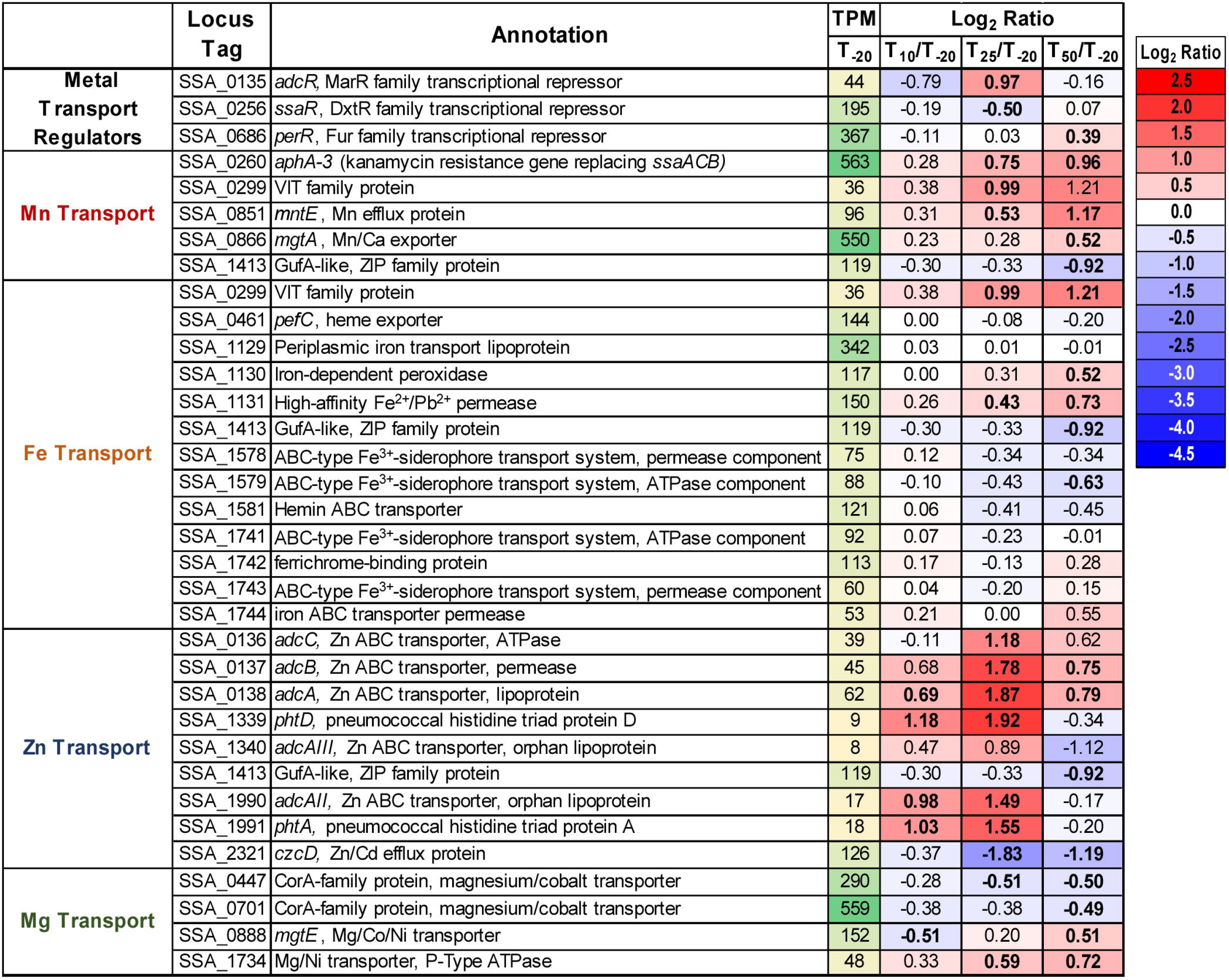
Expression of metal transport genes after Mn depletion. Putative metal transport genes are depicted with their average transcripts per million reads (TPM) at T_−20_ and log_2_ fold change values for each post-EDTA time point. TPM values greater than 1000 are full saturation (green). Positive log_2_ fold change values (red) indicate genes upregulated in after Mn depletion samples as compared to T_−20_, while negative values (blue) indicate downregulated genes. Values in bold indicate significant changes in expression by adjusted *P*-value (≤ 0.05).

Given that the cells were Mn depleted after EDTA addition (Fig 1), it was surprising that genes encoding putative orthologs of the *S. pneumoniae* Mn-export proteins MntE (Jakubovics and Valentine, 2009;Rosch et al., 2009;Martin and Giedroc, 2016) and MgtA (Martin et al., 2019), were significantly upregulated at T_50_ (Fig. 3). Expression of *mntE* was found to be constitutively expressed in *S. pneumoniae* (Lisher et al., 2013), and *mgtA* expression was found to be positively regulated by Mn through a metal-dependent riboswitch (Martin et al., 2019). We therefore sought to test whether previously generated Δ*mntE* and Δ*mgtA* mutants (Xu et al., 2011) exhibited increased Mn sensitivity relative to WT as expected based on previous findings in *S. pneumoniae* (Rosch et al., 2009;Martin et al., 2019). The Δ*mgtA* mutant grew as expected, with a lower final density than WT in BHI with 2 mM added Mn (data not shown). The Δ*mntE* mutant grew just like WT, which was not expected (data not shown). We tested up to 10 mM Mn in BHI and Todd Hewitt broth with 1% yeast extract and could not find a Mn concentration that prevented growth of either the Δ*mntE* mutant or WT (data not shown). We also generated an IPTG-inducible complemented mutant and did not observe a difference in growth in BHI with or without Mn upon addition of IPTG (data not shown). Initial metal analysis revealed that the Δ*mntE* mutant accumulated slightly more Mn than WT (data not shown). These results indicate that *S. sanguinis* may primarily use MgtA to export excess Mn and MntE may function differently in *S. sanguinis* than in *S. pneumoniae*. Future studies will elucidate the function of these putative exporters and their transcriptional regulation in *S. sanguinis*.

As seen in Fig. 1, cellular Zn levels in Δ*ssaACB* were not significantly altered by EDTA addition, despite the high affinity of this chelator for Zn (Perrin and Dempsey, 1974). Zn level maintenance may be due to the higher levels of Zn than Mn in BHI (1.7 ± 0.02 vs. 0.02 ± 0.003 μg ml^−1^, respectively) (Murgas et al., 2020) or the regulation of Zn transporter genes. In *S. pneumoniae*, transcription of the operon encoding the Zn ABC transporter AdcCBA (Dintilhac and Claverys, 1997) is repressed by a Zn-dependent, MarR-family regulator called AdcR (Shafeeq et al., 2011;Manzoor et al., 2015). *S. sanguinis* possesses orthologs of the *adcR* and *adcCBA* genes, and the latter genes were upregulated post-EDTA (Fig. 3). Expression of the gene encoding the Zn^2+^ and Cd^2+^ efflux protein, CzcD (Nies, 1992), was also down regulated after EDTA addition (Fig. 3). Thus, cellular Zn levels appear to have been maintained during EDTA treatment by decreasing export of intracellular Zn and increasing import of any remaining bioavailable Zn.

We also examined the regulation of other putative Zn-transport proteins. In *S. pneumoniae*, AdcAII is an orphan lipoprotein of the AdcCBA system (Bayle et al., 2011;Plumptre et al., 2014) encoded adjacent to a histidine triad protein also implicated in Zn transport (Bersch et al., 2013;Kallio et al., 2014). Interestingly, *S. sanguinis* has two genes, SSA_1340 and SSA_1990, that encode proteins similar to AdcAII, and each is also adjacent to putative histidine triad protein genes, SSA_1339 or SSA_1991. Because AdcAII is more similar to SSA_1990, we have named this protein AdcAII, whereas we have designated SSA_1340 AdcAIII. Consistent with a potential role in Zn uptake, all four of these genes were upregulated at T_25_ (Fig. 3). The relative contribution of each of these proteins to Zn import remains to be determined, although we hypothesize that the upregulation of these orphan lipoproteins and histidine triad proteins contributes to the tight maintenance of Zn levels in cells post-EDTA.

Less is known about the transport of other metals in streptococci. *S. sanguinis* encodes several putative Fe transporters, none of which were differentially expressed, with the exception of a vacuolar iron transporter (VIT) family homolog (Fig. 3). While this protein family has not been well-characterized in bacteria (Bhubhanil et al., 2014), VIT proteins have been implicated in Fe and Mn transport in other organisms (Li et al., 2001;Kim et al., 2006;Labarbuta et al., 2017). Two predicted CorA-family Mg transporters (Kehres et al., 1998;Groisman et al., 2013) were slightly downregulated post-EDTA (Fig. 3). This is unsurprising, as levels of Mg in BHI are very high (15.0 ± 1.5 μg ml^−1^) (Murgas et al., 2020), and EDTA has a lower affinity for Mg than many other metals (8.7 log_β1_ for Mg vs. 14.1 log_β1_ for Mn) (Perrin and Dempsey, 1974). For reasons that are unclear, expression of genes for two other putative Mg transporters, *mgtE* and *mgtB* (Groisman et al., 2013), was significantly upregulated post-EDTA (Fig. 3). The role and contribution of each of these gene products to metal homeostasis needs to be validated for *S. sanguinis*.

### 3.4 Examination of known Mn-cofactored enzymes

*S. sanguinis* possesses a singular superoxide dismutase, SodA, and it is Mn-cofactored (Xu et al., 2007;Crump et al., 2014). Our previous study indicated that reduced SodA activity could account for only a portion of the reduced virulence and serum growth of the Δ*ssaB* mutant (Crump et al., 2014). Expression of *sodA* decreased significantly at both T_25_ and T_50_ despite constant air input (Fig. 4), which may be due to Mn-dependent positive regulation of transcription (Jakubovics et al., 2002;Eijkelkamp et al., 2014). Given that the fermentor growth conditions do not exactly replicate either of our previous *in vitro* or *in vivo* assays, we wondered whether SodA would be important here. To answer this question, we grew our Δ*sodA* knockout mutant in the fermentor under the same conditions without EDTA. The Δ*sodA* mutant grew similarly to WT (Fig. S5), indicating that Mn-dependent SodA activity is not essential for aerobic growth under these conditions. While this does not rule out the possibility that reduced SodA activity after Mn depletion contributed to the reduced growth rate of Δ*ssaACB*, it established that it was not the sole cause, thus encouraging us to investigate other possibilities.

**Figure 4.**
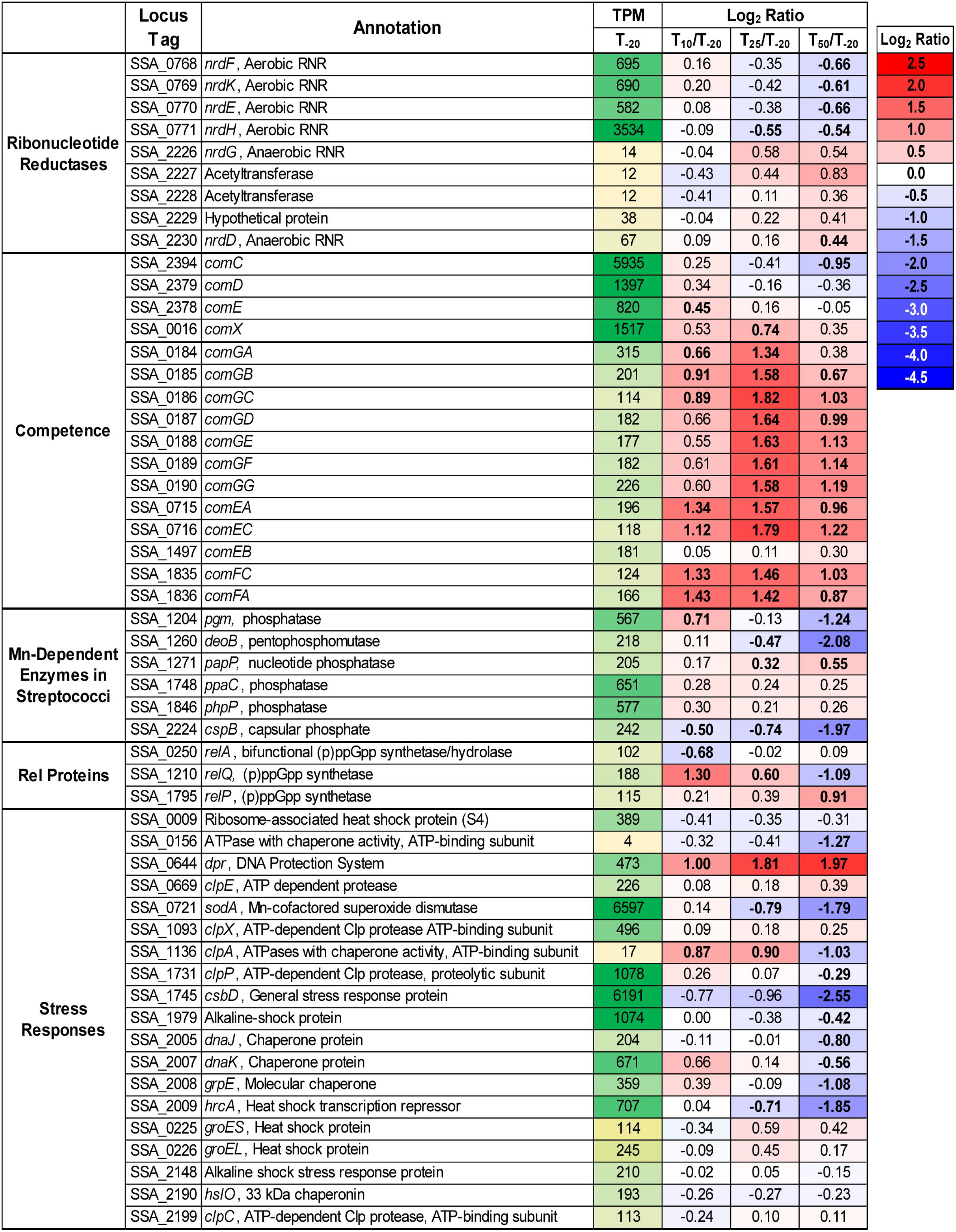
Expression of select genes after Mn depletion. Selected genes of interest are depicted with their average transcripts per million reads (TPM) at T_−20_ and log_2_ fold change values for each post-EDTA time point. TPM values greater than 1000 are full saturation (green). Positive log_2_ fold change values (red) indicate genes upregulated in after Mn depletion samples as compared to T_−20_, while negative values (blue) indicate downregulated genes. Values in bold indicate significant changes in expression by adjusted *P*-value (≤ 0.05).

The other known Mn-cofactored enzyme in *S. sanguinis* is the aerobic class Ib ribonucleotide reductase (RNR), NrdEF (Makhlynets et al., 2014;Rhodes et al., 2014). RNR enzymes catalyze the production of dNTPs from NTPs. It was previously found that mutant strains lacking this enzyme were unable to grow in aerobic conditions, whether in serum or BHI. These studies also suggested that Fe could not substitute for Mn as an RNR cofactor *in vivo*, despite its ability to do so *in vitro*. Thus, we considered whether loss of activity of the NrdEF enzyme due to Mn depletion was the cause of the observed growth rate decrease.

Expression of *nrdHEKF* was downregulated after Mn depletion (Fig. 4). In *Streptomyces coelicolor* (Borovok et al., 2004) and *Escherichia coli* (Torrents et al., 2007), the expression of ribonucleotide reductase genes is controlled by the NrdR repressor, which binds to NrdR boxes in response to increased dATP:ATP ratios, thus reducing expression when dNTPs are abundant (Grinberg et al., 2006). In SK36, putative NrdR boxes (Rodionov and Gelfand, 2005) were identified upstream of *nrdH* using RegPrecise (Novichkov et al., 2013) (https://enigma.lbl.gov/regprecise/; RRID:SCR_002149; data not shown). Thus, if regulation occurs by a similar mechanism in *S. sanguinis*, the decreased expression of the *nrdHEKF* operon (Fig. 4) would suggest that dNTPs were plentiful. We next considered whether the cells were able to obtain nucleotides from some other source. Although expression of the anaerobic RNR genes *nrdD* and *nrdG* slightly increased after Mn depletion, it seems highly unlikely that this could compensate for loss of NrdEF activity; the anaerobic enzyme is highly sensitive to oxygen in *S. sanguinis*, as in other bacteria (Rhodes et al., 2014). Moreover, the expression level of the *nrdG* gene at the T_−20_ time point was one-fortieth that of any of the genes in the *nrdHEKF* operon.

As *S. sanguinis* is naturally competent (Gaustad and Havarstein, 1997), we also considered the possibility that it was compensating for reduced NrdEF activity through the uptake of DNA from its environment. While several early competence genes (Rodriguez et al., 2011) were either unchanged or downregulated, *comX* and most of the late competence genes were upregulated significantly at T_25_ (Fig. 4). Interestingly, this upregulation was sustained at T_50_ for most genes, despite the fact that competence has been characterized as a transient state in *S. sanguinis* (Rodriguez et al., 2011). Elimination of genetic competence genes *comCDE* or *comX* (Callahan et al., 2011) did not influence aerobic growth in Mn-deplete media in the WT or Δ*ssaACB* background (data not shown). We recently analyzed the metabolome of *S. sanguinis* cells under the same conditions as this study (unpublished data). The results suggested that deoxynucleotides in cells increased after Mn depletion, which is inconsistent with a deoxynucleotide shortage. Thus, while NrdEF requires Mn for activity, our data suggest that nucleotides were not a limiting factor for growth in our study.

The simplest explanation for the above results is that *S. sanguinis* possesses at least one other Mn-dependent, essential enzyme in addition to NrdEF, and when Mn levels fall, the reduced activity of one or more of these other enzymes becomes growth limiting. In the related species *S. pneumoniae*, there are six additional enzymes (Fig. 4) that have been found to be co-factored by Mn (Kuipers et al., 2016;Martin et al., 2017). Orthologs of all six enzymes are encoded in the *S. sanguinis* genome, although their functions have not been confirmed. Pgm and CpsB are phosphatases that have been implicated in capsule biosynthesis in *S. pneumoniae,* and both of these genes were significantly downregulated at T_50_. DeoB is a phosphopentomutase that functions to connect the pentose phosphate pathway to purine biosynthesis and was also significantly downregulated at T_50_. Expression of *papP*, encoding a nucleotide phosphatase, was significantly increased at the later time points and has been shown to affect membrane lipid homeostasis (Kuipers et al., 2016). A significant morphological difference was observed in Δ*papP* mutants in *S. pneumoniae,* but Δ*ssaACB* cells from the T_50_ sample did not appear morphologically different from cells at T_−20_ (data not shown). Of note, we observed changes in fatty acid synthesis under these same fermentor growth conditions (unpublished data), suggesting that PapP activity may be reduced but not to the extent required to affect morphology. Genes encoding the other two phosphatases, PhpP and PpaC, were not differently expressed at any time point (Fig. 4). While this indicates the lack of a Mn-dependent regulation mechanism, it does not rule out the possibility that their activity decreased. PhpP is a serine/threonine protein phosphatase that is a key regulator of cell division and has been shown to be regulated by the bioavailable Zn:Mn ratio in *S. pneumoniae* (Martin et al., 2017). While the Zn:Mn ratio did increase over time in our study (Table 1), Δ*ssaACB* cells from the T_50_ sample did not appear morphologically different from cells at T_−20_ (data not shown), indicating that PhpP may not be affected by Mn limitation under these conditions. In our recent Tn-Seq study, loss of PhpP did not significantly affect the growth of *S. sanguinis* in serum (Zhu et al., *submitted*), which indicates that it is likely not responsible for the growth rate decrease observed here. The last phosphatase, PpaC, is essential for *S. sanguinis* (Xu et al., 2011), so if PpaC activity was decreased due to Mn depletion, this could have contributed to the decreased growth rate phenotype observed post-EDTA. Further studies utilizing the knockout mutants of each nonessential phosphatase (Xu et al., 2011) or an approach such as CRISPR interference (Shields et al., 2020) for PpaC will enhance our understanding of relative contributions of each phosphatase to the growth and morphology of *S. sanguinis*.

**Table 1.** Zn:Mn Ratios in Fermentor Grown Cells.

| Strain | T<sub>-20</sub> | T<sub>10</sub> | T<sub>25</sub> | T<sub>50</sub> |
| --- | --- | --- | --- | --- |
| WT | 1.64 | 1.66 | 1.75 | 1.81 |
| Δ*ssaACB* | 2.02 | 2.47* | 3.30* | 5.36* |

Values indicate ratios of Zn concentrations as compared to those of Mn. Concentrations were determined by ICP-OES and significance was determined by one-way ANOVA with a Tukey-Kramer multiple comparisons test for T<sub>-20</sub> where \* indicates $P \leq 0.05$ .

In streptococci and enterococci, Mn acts as a cofactor for the hydrolase domain of the bifunctional (p)ppGpp synthetase/hydrolase, RelA (also called RSH for <u>R</u>elA/<u>S</u>poT <u>H</u>omologs) (Mechold et al., 1996). As an alarmone, (p)ppGpp serves as an effector of the stringent response in bacteria (Irving and Corrigan, 2018). Expression of *relA* was unchanged after EDTA addition, and expression of the other two small alarmone synthetase genes, *relP* and *relQ* (Lemos et al., 2007), were significantly increased and decreased, respectively (Fig. 4). Both RelP and RelQ were found to produce less (p)ppGpp than RelA in *S. mutans* (Lemos et al., 2007) and appear to be important during different environmental conditions or growth stages in gram-positive bacteria (Yang et al., 2019).

In an attempt to determine whether loss of hydrolase activity in RelA could account for the phenotypes we observed in Mn-depleted cells, we attempted to construct a hydrolase-deficient mutant by altering specific residues (R44, H62, T151) shown by Hogg et al. (2004) to be important for (p)ppGpp hydrolase, but not synthetase activity. Similar to Kaspar et al. (2016), we were unable to generate any of the three point mutants without unintended mutations arising in other regions of the gene (data not shown). This indicates that hydrolase activity may be essential for growth of *S. sanguinis*. We then obtained strains from the comprehensive *S. sanguinis* mutant knockout library created by Xu et al. (2011) that were deleted for each of the *rel* genes. We also generated a *rel*^0^ strain by knocking out all three *rel* genes utilizing a markerless mutagenesis system originally described by Xie et al. (2011), but modified to contain the in-frame deletion cassette (IFDC) specific to *S. sanguinis* (Cheng et al., 2018). We also made the *rel* knockout mutants in the Δ*ssaACB* background. We then assessed the growth of these mutants in aerobic serum: our *in vitro* model for infective endocarditis (Crump et al., 2014). As shown in Fig. 5, neither Δ*relP* nor Δ*relQ* grew to a density that differed significantly from WT, whereas the growth of Δ*relA* was severely attenuated. Interestingly, the *rel*^0^ strain grew to a density that was significantly greater than Δ*relA* after 24 h, although not as great as WT. The Δ*ssaACB* parent strain grew poorly, as expected, and neither Δ*ssaACB* Δ*relP* nor Δ*ssaACB* Δ*relQ* grew to a density that was significantly different from the Δ*ssaACB* parent strain. The Δ*ssaACB* Δ*relA* strain grew significantly worse than the parent strain, and similar to the WT background, the Δ*ssaACB rel*^0^ mutant grew better than Δ*ssaACB* Δ*relA*.

**Figure 5.**
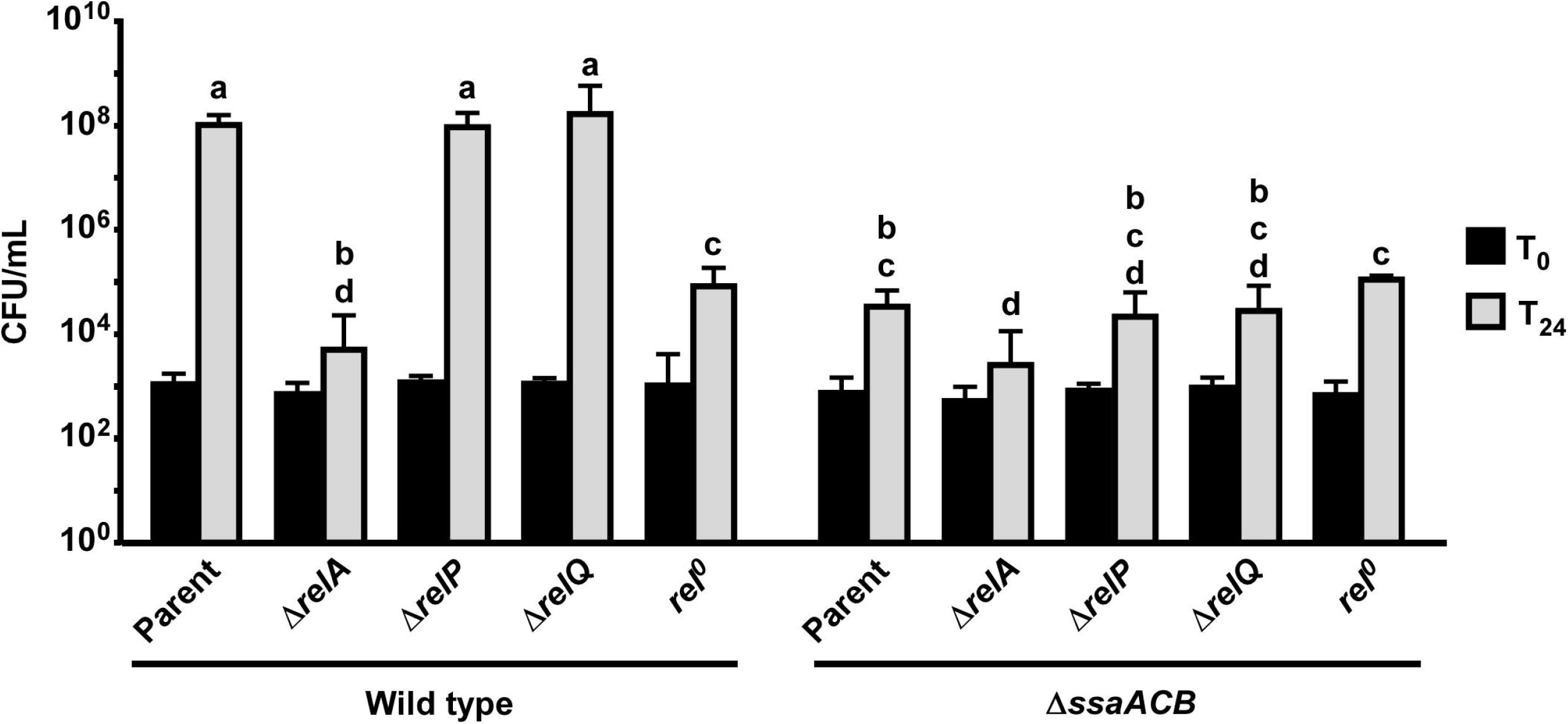
Aerobic serum growth of *rel* mutants. Various *rel* mutants were grown for 24 h in 6% O_2_ in pooled rabbit serum. The means and standard deviations of at least three replicates are displayed. Significance was determined by one-way ANOVA with a Tukey-Kramer multiple comparisons test. T_24_ bars with the same letter are not significantly different from each other. T_0_ bars from each strain were compared to each other by one-way ANOVA and found to be not significantly different.

In analyzing previous transcriptome studies in related species using Δ*relA* or Δ*rel*^0^ mutants producing little to no (p)ppGpp, we noted many expression patterns similar to our study (Nascimento et al., 2008;Kazmierczak et al., 2009;Colomer-Winter et al., 2019). While these studies utilized different species and growth conditions and thus are not a direct comparison to ours, it is remarkable that reduction in (p)ppGpp levels would lead to similar changes in gene expression as Mn depletion. Of special interest to us, several PTS were downregulated in all three previous studies. Similar to the results we observed in this study, the *S. pneumoniae* Δ*relA* mutant showed decreased expression of *spxB* and *sodA* (Kazmierczak et al., 2009). These comparisons indicate either that dysregulation of (p)ppGpp levels leads to changes in expression of these genes in response to stress, or that decreased activity of the Rel hydrolase domain is not responsible for the observed changes in expression of these genes.

Based on these results, we hypothesize that reduced activity of the RelA hydrolase domain may contribute to the observed reduction in growth rate in the fermentor studies but is not entirely responsible. Specifically, our inability to eliminate the cell’s only known (p)ppGpp hydrolase, combined with our finding that the Δ*relA* strains, which also have no hydrolase, exhibited worse growth than *rel*^0^ mutants having no synthetase and therefore no (p)ppGpp, suggests that (p)ppGpp accumulation is highly detrimental to growth. A definitive test of this hypothesis will require measurement of (p)ppGpp levels in fermentor-grown cells. We are currently assessing various approaches for feasibility. In addition, the significant decrease in growth of the Δ*ssaACB* Δ*relA* strain as compared to the Δ*ssaACB* parent shows that there is an additive effect, indicating that the impact of the loss of RelA is not entirely Mn-dependent.

### 3.5 Assessment of stress and stress responses in Mn-depleted cells through gene expression

We next sought to determine whether the RNA-seq data suggested anything concerning stresses experienced by the cells*. S. sanguinis* is known to generate copious amounts of hydrogen peroxide (H_2_O_2_), presumably to more effectively compete against other oral species, such as the caries-forming pathogen *S. mutans* (Kreth et al., 2005;Chen et al., 2011). Simple Mn compounds have been reported to prevent oxidative stress by catalyzing the decomposition of H_2_O_2_ (Liochev and Fridovich, 2004) and superoxide (Barnese et al., 2008;Barnese et al., 2012). We observed a significant decrease in expression of the gene encoding the H_2_O_2_-generating enzyme pyruvate oxidase, *spxB*, at T_25_ and T_50_ (Fig. 6A) (Spellerberg et al., 1996;Chen et al., 2011). To determine whether the decreased growth rate of the Δ*ssaACB* strain during aerobic fermentor growth after Mn depletion was due to excess H_2_O_2_ generation or the inability of cells to cope with H_2_O_2_ without Mn, H_2_O_2_ levels were measured in spent supernatant. Concentrations ranged between 1 and 5 μM (Fig. 6B), far lower than has been observed in previous studies employing SK36 (Kreth et al., 2008) despite the constant influx of air into the vessel. H_2_O_2_ levels also decreased significantly at T_25_ and T_50_ as compared to T_−20_ (Fig. 6B), which correlates with the decreased expression of *spxB* (Fig. 6A). These results indicate that oxidative stress related to excess H_2_O_2_ levels is unlikely to be the cause of the growth rate decrease observed after Mn depletion.

**Figure 6.**
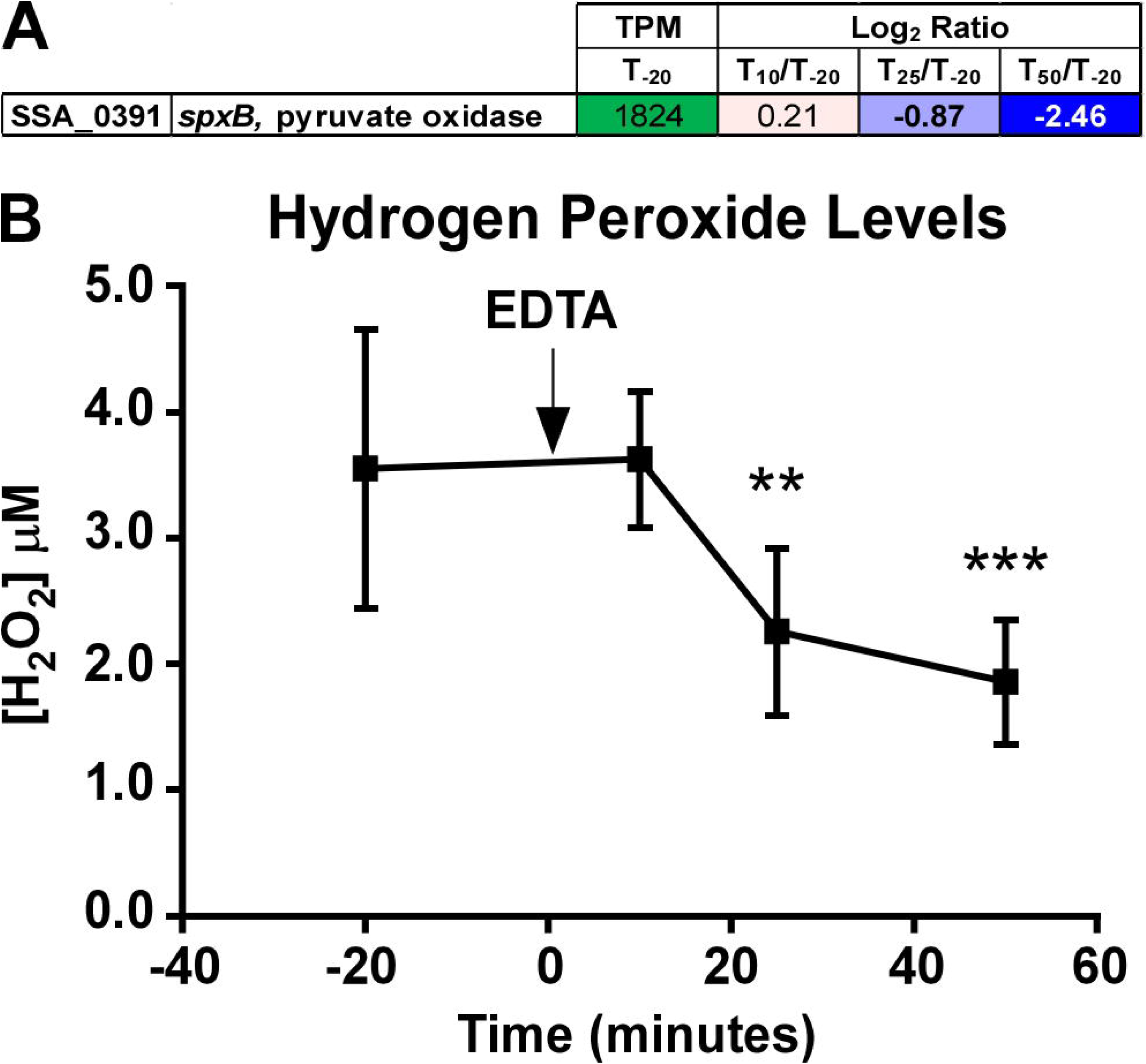
Quantitation of *spxB* expression and hydrogen peroxide in Δ*ssaACB* fermentor cultures. (**A**) Expression of the *spxB* gene in fermentor-grown Δ*ssaACB* cells as determined by RNA-seq analysis. Average transcripts per million reads (TPM) at T_−20_ and log_2_ fold change values for each after Mn depletion time point are displayed. TPM values greater than 1000 are full saturation (green). Positive log_2_ fold change values (red) are genes upregulated in after Mn depletion samples as compared to T_−20_, while negative values (blue) indicate downregulated genes. Values in bold indicate significant changes in expression by adjusted *P*-value (≤ 0.05). (**B**) H_2_O_2_ levels of the BHI culture supernatant were measured at each time point. Means and standard deviations of at least 4 replicates are shown. Significance was determined by one-way ANOVA with a Tukey-Kramer multiple comparisons test, comparing each after Mn depletion time point to T_−20_. ** *P* < 0.01, *** *P* <0.001.

Expression levels of various stress response genes were assessed, and most were either downregulated or unchanged at T_50_ (Fig. 4), indicating that the reduced growth rate is likely not due to an overwhelming stress response. The only stress response-related gene to show a significant increase in expression at T_50_ was that encoding the Dps-like peroxide resistance protein, Dpr, (Yamamoto et al., 2000). Dpr is a ferritin-binding protein that has been shown to be imperative for oxidative stress tolerance in several streptococci (Yamamoto et al., 2000;Yamamoto et al., 2002;Pulliainen et al., 2003;Brenot et al., 2005), including *S. sanguinis* SK36 (Xu et al., 2014) and was one of the most highly upregulated genes at all three time points. During initial studies of a Δ*dpr* mutant created previously (Xu et al., 2011), as well as a strain with this mutation in a Δ*ssaACB* background, each strain grew slowly in BHI under the same aerobic fermentor conditions used in this study (data not shown). These results indicate that Dpr may contribute to aerobic growth in a Mn-independent manner and increased expression after Mn depletion may be due to augmented oxidative stress.

### 3.6 Analysis of carbon catabolite repression and sugar transport

Examination of transport gene clusters revealed that the majority of those thought to transport sugars were downregulated (Table S2), and of these, the majority belonged to the PTS family, which is regulated by carbon catabolite repression (CCR). CCR is a regulatory mechanism that gives bacteria the ability to utilize carbon sources in order of preference (Gorke and Stulke, 2008). In gram-positive bacteria, a carbon catabolite protein such as CcpA binds to catabolite responsive elements (*cre*) and represses transcription of genes encoding non-preferred carbon source transport and utilization systems (Warner and Lolkema, 2003). To determine the extent to which CcpA binding could be responsible for the observed downregulation, *cre* sites identified previously by RegPrecise (Novichkov et al., 2013) and by our custom searches were collected and compared. Using these methods, 393 putative binding sites were identified (Table S2). Several PTS and sugar ABC transport genes were predicted to have 5’ *cre* sites, the majority of which were downregulated at T_50_. Other genes known to be CcpA-regulated, such as *spxB* (Zheng et al., 2011;Redanz et al., 2018b), were downregulated as well. This is surprising given that the glucose-containing media was replenished at a constant and rapid rate throughout the experiment, indicating that there could be a Mn-related mechanism for CcpA repression. Cells also did not appear to be starved for glucose; when excess glucose (1.8%) was added to the media, the post-EDTA growth rate was similar to that of normal BHI, which contains 0.2% glucose (data not shown). This is not entirely unexpected, as it has been established by Redanz et al. (2018b) that CcpA repression of *spxB* in *S. sanguinis* is glucose-independent. One possible explanation for the Mn-dependent, glucose-independent CcpA repression observed in our study is the accumulation of the glycolytic intermediate, fructose-1,6-bisphosphate (FBP) during Mn depletion (Fig. 7). In Firmicutes, phosphorylation of histidine phosphocarrier protein (HPr) to HPr(Ser-P) occurs when FBP and ATP levels are high (Gorke and Stulke, 2008). HPr(Ser-P) then binds to CcpA, which in turn induces the binding of the repressor to *cre* sites on the DNA. Additionally, FBP enhances the binding interaction of HPr(Ser-P) and CcpA, increasing repression. In our concurrent metabolomics study, we found that Δ*ssaACB* cells accumulated high levels of FBP at T_50_ after Mn depletion (unpublished data), which may explain the strong evidence for CcpA repression. As expected, if CcpA were responsible for the changes in expression, we found that of the 169 DEGs found by (Bai et al., 2019) when comparing a *ccpA* mutant to SK36, 48 were changed in the opposite direction as our T_50_ sample. However, 15 significant DEGs were in the same direction, and the remainder were unchanged in our study. Additionally, most of the DEGs we observed in our study did not overlap with those of Bai et al. (2019). This comparison indicates that CcpA-dependent repression could be responsible for some of the changes in expression after Mn depletion, but it does not explain all of the observed results.

**Figure 7.**
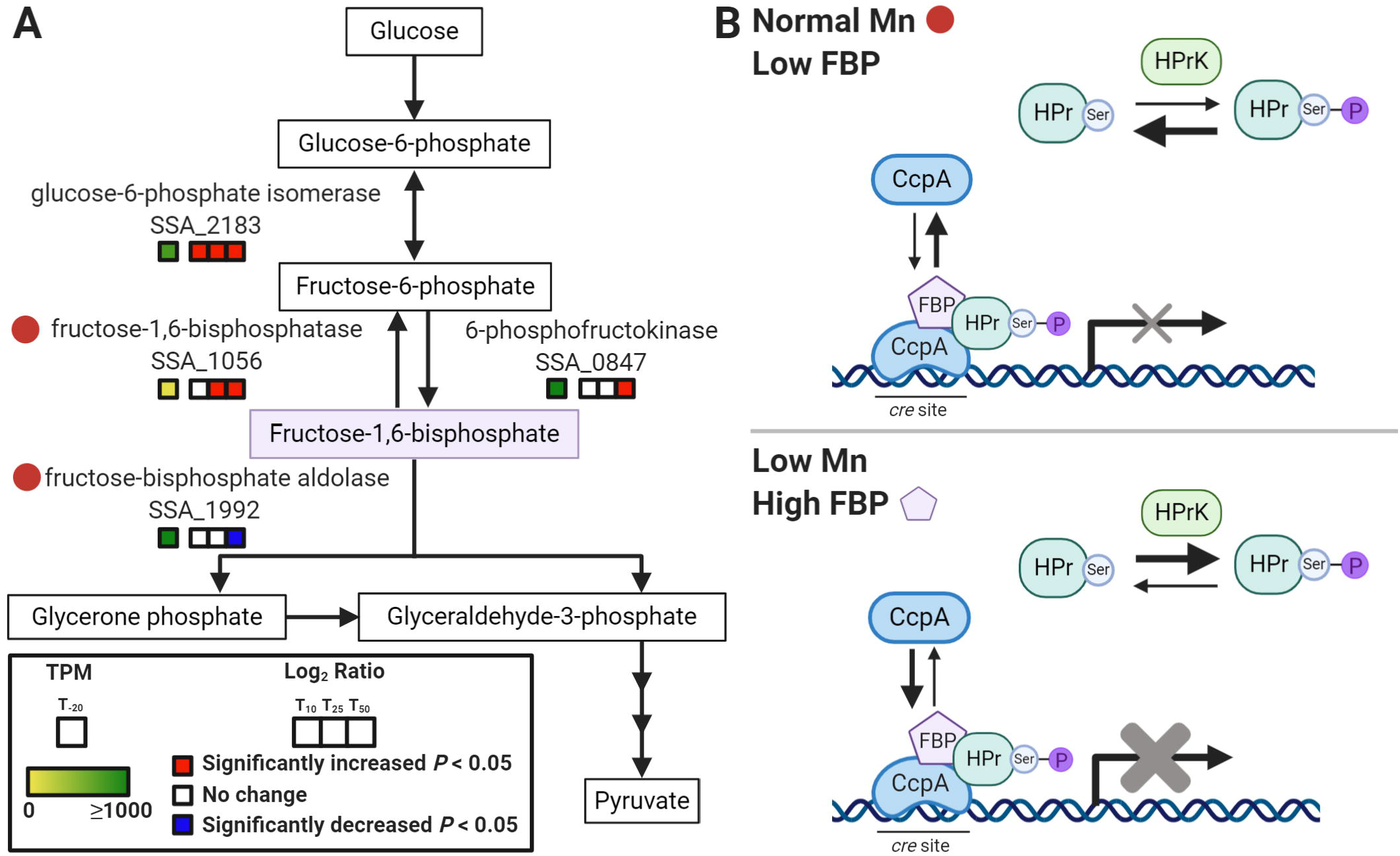
Model of Mn-dependent CcpA repression. (**A**) Depiction of a segment of the glycolysis and gluconeogenesis pathway in *S. sanguinis* from KEGG (Kanehisa and Goto, 2000). Red circles indicate potentially Mn-cofactored enzymes. Gene expression for each post-EDTA time point compared to T_−20_ is indicated by the colored boxes. Significant changes (*P* < 0.05) are indicated by red (increased) or blue (decreased). (**B**) Model of Mn-dependent CcpA repression based on CCR in Firmicutes as described by Gorke and Stulke (2008). In the top panel, normal Mn levels in BHI result in low FBP, which leads to less phosphorylation of HPr to HPr(Ser-P) by HPrK. With low FBP and HPr(Ser-P), CcpA exists mainly in its free state, unbound to *cre* sites in the DNA. This results in little to no repression of the CcpA regulon. The bottom panel depicts Mn depletion, where reduced activity of fructose-1,6-bisphosphatase and fructose-bisphosphate aldolase leads to an accumulation of FBP. This induces the phosphorylation of HPr so that it is primarily in the HPr(Ser-P) state. Increased FBP levels also enhance binding of CcpA to HPr(Ser-P) and to DNA. This results in augmented repression of the CcpA regulon.

### 3.7 Other findings

We observed that several amino acid transporters and synthetases were differentially regulated after Mn depletion (Fig. S6A). We were interested in learning whether a defect in amino acid synthesis could be responsible for the observed growth rate reduction. Addition of casamino acids resulted in slightly improved growth of Δ*ssaACB* cells in 12% O_2_ and pooled rabbit serum (data not shown), an *in vitro* environment that mimics the conditions of the aortic heart valve (Crump et al., 2014). Further studies were conducted wherein individual amino acids were added alone or in addition to casamino acids, and it was determined that the addition of 4 μM cysteine alone improved the growth to WT-like levels (Fig. S6B). Cysteine also improved growth of Δ*ssaACB* cells in BHI + 100 μM EDTA in static cultures set to 12% O_2_, although not quite to WT-like levels (Fig. S6C). Addition of the reduced form of the tripeptide glutathione (γ-L-glutamyl-L-cysteinyl-glycine; GSH) led to similar results (Fig. S6). Both cysteine and GSH can act as antioxidants (Potter et al., 2012), which may explain the improvement in aerobic tube growth. Interestingly, though, neither improved the growth of Δ*ssaACB* cells cultured in the fermentor with EDTA (data not shown). We hypothesize that this difference may be due to the increased aeration in the fermentor vessel from the constant influx of air as compared to the static tube cultures.

Finally, we also observed decreased expression of large, contiguous loci encoding ethanolamine utilization, a type IV pilus system, and CRISPR-associated proteins (Fig. S7), in addition to many smaller loci and individual genes (Table S1).

## 4 Discussion

Only two enzymes have been confirmed to be Mn-dependent in *S. sanguinis,* and few others have been identified in other streptococci. Despite this, we observed changes in a wide variety of systems after Mn depletion of the Δ*ssaACB* mutant using EDTA. One possible explanation for this discrepancy is that Mn binds with low affinity to most proteins, resulting in Mn loss or replacement during purification. In fact, initial studies of the aerobic class Ib RNR identified Fe as the exclusive cofactor based on RNR activity *in vitro* and the fact that Fe was present in many different bacterial RNRs heterologously expressed in *E. coli*. Only later was it discovered that these enzymes were Mn-cofactored when natively expressed (Cotruvo and Stubbe, 2012), and despite the *in vitro* activity of both forms of the *S. sanguinis* RNR, only the Mn-cofactored version was active *in vivo* (Makhlynets et al., 2014;Rhodes et al., 2014). Additional Mn-dependent enzymes may have similarly escaped detection. Another possible explanation is that Mn depletion impacts several key regulatory systems, such as CCR and (p)ppGpp, which leads to changes in the expression of many different genes. Mn levels have been found to be related to each of these systems in other gram-positive bacteria (Kehres and Maguire, 2003;Colomer-Winter et al., 2017). Here we highlight Mn-related systems we identified in this study of *S. sanguinis* for future investigation.

Given that BHI contains glucose, it was expected that *S. sanguinis* would preferentially transport and utilize it as a preferred carbon source under standard fermentor conditions. This is supported by the fact that glucose levels decreased in the media after cell growth in our corresponding study (unpublished data) as well as by the high expression of putative glucose transporters SSA_1752, SSA_1918-1920, and SSA_1298-1300 (Ajdic and Pham, 2007) at T_−20_ (Table S2). Surprisingly, expression of nearly all sugar transport systems decreased after Mn depletion (Table S2), despite nearly constant levels of glucose in the cells (unpublished data). CcpA is known to repress its own expression in a glucose-dependent manner (Bai et al., 2019), and yet much like the glucose transporters, *ccpA* expression was high at T_−20_ and significantly decreased by T_50_ (Table S2). Potential explanations could include: (*i*) 2 g/L glucose in BHI is not sufficient to induce CcpA repression; (*ii*) other regulatory mechanisms are preventing proper CCR under these conditions; or (*iii*) much like *spxB*, many other systems in *S. sanguinis* are subject to glucose-independent CcpA repression. Redanz et al. (2018a) used 0.3% as the low glucose condition in their study of CcpA-repression of *spxB*, whereas Bai et al. (2019) used BHI alone (0.2% glucose) to observe differences in the transcriptome between the WT and Δ*ccpA* strains. Thus, the glucose concentration of BHI may indeed be low, yet sufficient to induce some repression of its regulon.

Additionally, it is unexpected that Mn depletion leads to an apparent increase in CcpA repression. The strongest evidence for CcpA-dependent regulation is *spxB.* In *S. sanguinis*, *spxB* expression has been shown to be positively regulated by SpxA1 (Chen et al., 2012) and VicK (Moraes et al., 2014) and negatively regulated by CcpA (Zheng et al., 2011). The *spxA1* gene was in the top 10% of all genes based on expression at T_−20_ and remained unchanged after EDTA addition (Table S1), indicating that repression by CcpA is likely responsible for the decrease in *spxB* expression as opposed to changes in induction by SpxA1. The mechanism by which CcpA represses *spxB* expression in *S. sanguinis* is unique from other streptococci in that it is independent of glucose (Redanz et al., 2018b). It was previously determined that Mn may play a role in *spxB* expression in *S. pneumoniae*, as a Δ*mntE* mutant in *S. pneumoniae* accumulated Mn and produced more H_2_O_2_ than WT under excess Mn conditions (Rosch et al., 2009).

The connection between Mn and sugar catabolism is not unprecedented, as previous studies have implicated Mn as important for the activity of sugar catabolism enzymes in other bacteria (Kehres and Maguire, 2003). The accumulation of FBP might explain the glucose-independent CcpA repression we observed in most genes with putative *cre* sites (Table S2). To determine potential causes for the accumulation, we examined the enzymes required for synthesizing and catabolizing FBP. We noted that in UniProt (uniprot.org; RRID:SCR_004426), fructose-1,6-bisphosphatase (Fbp; SSA_1056) is annotated as Mn-cofactored. Fbp is the only enzyme in *S. sanguinis* known to catalyze the reaction from FBP to fructose-6-phosphate (F6P) through gluconeogenesis, as depicted in KEGG (https://www.genome.jp/kegg/; RRID:SCR_012773) (Kanehisa and Goto, 2000). While this may contribute to an accumulation of FBP, it is unlikely to be the principal factor, as expression levels of *fbp* are low at T_−20_ (Fig. 7A). Another contributing factor may be fructose-bisphosphate aldolase (Fba; SSA_1992), which catalyzes the production of glyceraldehyde 3-phosphate and glycerone phosphate from FBP. While the UniProt annotation states that SSA_1992 is co-factored by Zn, BRENDA (brenda-enzymes.org; RRID:SCR_002997)(Jeske et al., 2019) shows that several Fba orthologs are cofactored by Mn, including one from *Deinococcus radiodurans* (Zhang et al., 2006). Thus, it is possible that reduced activity of both Fba and Fbp due to Mn depletion led to the accumulation of FBP, which in turn induced CcpA repression after Mn depletion (Fig. 7). Enzymatic activity assays will be required to determine the true cofactor for these enzymes in *S. sanguinis*, but accumulation of FBP is strong evidence that the activity of at least one enzyme in this pathway is affected by Mn depletion.

In addition to repression of sugar transport related genes, we also identified a multitude of other repressed genes that we either confirmed were members of the CcpA regulon (Bai et al., 2019) or that had putative *cre* sites (Table S2). Three of these genes are annotated as Mn-cofactored phosphatases in *S. pneumoniae*. The gene *pgm* has putative 5’ *S. suis cre* and *cre*2 sites; the *cpsA* gene, immediately upstream of *cpsB,* has a putative *cre*2 site; and *deoB* is encoded in an operon with *rpiA* and *punA*, confirmed members of CcpA regulon (Bai et al., 2019). Pgm and CpsB (Morona et al., 2002;Tsunashima et al., 2012;Skov Sørensen et al., 2016) are implicated in capsule formation, although *S. sanguinis* lacks a true capsule. As far as we know, “capsule” formation genes have not been implicated in cell growth, and decreased Mn levels due to loss of SsaB was not found to impact colonization of the sterile heart valve vegetation after 2.5 h in a rabbit model of IE (Crump et al., 2014). DeoB links glucose metabolism with purine biosynthesis (Tozzi et al., 2006;Panosian et al., 2011). As mentioned above, nucleotide levels seem to be unaffected as evidenced by lack of NrdR repression of *nrdF* (Fig. 4). Thus the impact of decreased *deoB* expression does not appear to affect nucleotide levels, although it may have other unappreciated effects on the cell. The connection between these enzymes and CcpA provides further support for the theory that Mn levels are related to CCR.

The relationship between (p)ppGpp and carbon source utilization has been previously established; *S. mutans* strains lacking RelA or all Rel proteins showed delayed growth rates when transitioned from media containing glucose to lactose (Zeng et al., 2018). When we examined the genome for *cre* sites, *relQ* was predicted to have a 5’ *S. suis cre* site and was downregulated at T_50_ (Table S2), indicating that it could be under CcpA control. Very little is known about the transcriptional regulation of *relA* in streptococci (Nascimento et al., 2008;Irving and Corrigan, 2018), although regulation of activity has been established in other species (Gratani et al., 2018). Expression of *relP* and *relQ* appears to be growth phase-dependent, as well as regulated by environmental factors (Ronneau and Hallez, 2019). In *S. mutans*, expression of *relP* is activated by a two-component system, RelRS, which is thought to sense oxidative stressors (Seaton et al., 2011). It was hypothesized by Kim et al. (2012) that (p)ppGpp production by RelP in *S. mutans* may be an attempt by the cell to slow growth to minimize damage from oxygen radicals produced during metabolism. While it was observed in this study that H_2_O_2_ levels decreased in response to EDTA addition, it is possible that other ROS were present due to a decrease in SodA activity. Thus, increased expression and activity of RelP, in addition to lack of hydrolase activity by RelA, could be at least partly responsible for the reduced growth rate.

Similar to our study, metals and CCR have been found to impact amino acid transport in other gram-positive bacteria. In *B. subtilis,* expression of biosynthetic genes for amino acids such as arginine, cysteine, and histidine was affected by addition of excess metals (Moore et al., 2005). In addition, expression of amino acid transporters and synthetases has been shown to be regulated by various CCR mechanisms (Jourlin-Castelli et al., 2000;Bai et al., 2019), as well as by VicRK orthologs (Radin et al., 2016). In *Staphylococcus aureus*, replacing sugars with amino acids for glycolysis reduced the need for cellular Mn (Radin et al., 2016;Parraga Solorzano et al., 2019). VicRK may also be responsive to Mn through SsaR, as the *S. mutans* ortholog, SloR, was shown to regulate VicRK expression (Downey et al., 2014). Consistent with this, the expression of VicR (SSA_1565) was slightly, yet significantly decreased at T_50_ (Table S1). Thus, our results implicate Mn depletion as another potential influence on the regulation of amino acid transport and synthesis, either directly by decrease in function of an unidentified Mn-dependent enzyme or indirectly by influencing other regulatory systems, such as CCR or VicRK.

In this study, all of the ethanolamine utilization (*eut*) genes were downregulated at T_25_ and T_50_ (Fig. S6). Ethanolamine (EA) is a potential carbon and nitrogen source for bacteria derived from phosphotidylethanolamine found in membranes (Kaval and Garsin, 2018). Much like the gut microbe *Enterococcus faecalis* (Li et al., 2017), the *S. sanguinis* genome (Xu et al., 2007) encodes a large cluster of *eut* and 1,2-propanediol utilization (*pdu*) genes (Kaval et al., 2019). In *E. faecalis*, *eut* expression is positively regulated in the presence of EA by a two-component system (TCS) composed of the proteins EutW and EutV (Fox et al., 2009). Recently, it was discovered that the EutV/EutW TCS itself is negatively regulated by CcpA, and there are multiple putative *cre* sites within the *eut* gene cluster (Kaval et al., 2019). These *cre* sites were not found in the RegPrecise search of *S. sanguinis* (Table S2), which was also noted by Bai et al. (2019). The exact role of EA catabolism in *S. sanguinis* growth is still to be determined. Of note, despite a putative *cre* site upstream of *pduB* (Kaval et al., 2019), expression of the *pdu* gene cluster was unchanged in our study (Table S1). However, this may be due to minimal expression pre-EDTA (T_−20_).

Expression of most clustered regularly interspaced short palindromic repeats (CRISPR)-associated proteins was significantly decreased in the latter two post-EDTA time points (Fig. S6). CRISPR-associated proteins that make up the CRISPR-Cas system are found in 40% of bacterial species (Makarova et al., 2015). In addition to providing cells with adaptive immunity against foreign mobile elements, including phages and plasmids, these systems have been implicated in modulating oral biofilm development, DNA repair, and DNA uptake (Gong et al., 2020). Of particular interest, the type I and type II CRISPR systems of *S. mutans* have also been implicated in modulating stress response phenotypes, such as to pH, temperature, and oxidative stress (Serbanescu et al., 2015). The type I mutant was able to grow faster in an acidic environment but was more sensitive to H_2_O_2_, paraquat, and SDS (Serbanescu et al., 2015). Oral bacteria tend to have either type I or type II systems (Gong et al., 2020) but *S. sanguinis* SK36 has a type III system as determined by CRISPRCasFinder (Couvin et al., 2018).

The decrease in expression was unexpected, as the Δ*ssaACB* fermentor-grown cells are likely experiencing more oxidative stress due to decreased expression and function of SodA. Thus, it is possible that there is another function or mechanism for regulation of the CRISPR-associated protein genes. Additionally, Csm1 (SSA_1251) was found to be a part of the CcpA regulon (Bai et al., 2019), although no *cre* site was identified (Table S2). The role and regulation of the CRISPR-Cas system of *S. sanguinis* SK36, as well as its relationship to Mn levels, should be examined further.

The expression of a type IV pilus was significantly downregulated after Mn depletion (Fig. S6). *S. sanguinis* is unusual in that it is the only streptococcal species to encode a gene cluster for the biosynthesis of a type IV pilus system (T4P) (Gurung et al., 2016) that is distinct from the T4P competence pilus (Sheppard et al., 2020), and yet few strains exhibit the twitching motility normally mediated by T4P (Chen et al., 2019). In SK36, T4P appears to be important for adherence to host cells and may be regulated by CcpA, despite the lack of a *cre* site in the promoter region (Bai et al., 2019;Chen et al., 2019). In *S. pneumoniae*, the SsaR ortholog, PsaR, negatively regulates the expression of type I pilus genes in the presence of Mn (Johnston et al., 2006) whereas a Δ*mntE* mutant containing excess intracellular Mn showed increased expression of the same genes (Rosch et al., 2009). While the pilus type differs between these two streptococcal species, the shared Mn-dependent regulation may indicate that interplay between the Mn homeostasis and pilus expression is a conserved characteristic. Further studies are required to investigate this relationship and its role in the virulence of *S. sanguinis*.

The effect of Mn depletion on a multitude of diverse systems indicates that the impact of Mn is not relegated to only a few enzymes. Depletion of Mn does not induce a traditional stress response, instead inducing what appears to be dysregulation of many different genes that leads to rapid reduction in the growth rate, despite plentiful nutrients and other metals. While decreased function of the known Mn-cofactored enzymes, such as NrdF, SodA, and the hydrolase domain of RelA, likely contributed to the decreased growth rate we observed upon Mn depletion, it is probably a combination of multiple systems leading to the observed phenotype. Additionally, a majority of the affected systems appear to be regulated by CCR through CcpA-dependent repression in a glucose-independent manner. Future research will focus on determining the respective contribution of each putative Mn-dependent enzyme as well as whether there is a direct relationship between Mn and CCR.

## 5 Materials and Methods

### 5.1 Bacterial strains and growth conditions

The *S. sanguinis* strain SK36 is a human oral isolate from Mogens Killian, Aarhus University, Denmark. All mutant strains were generated in the SK36 background (Table 2). For the Δ*ssaACB* strains, all three genes were replaced with either a kanamycin (Kan) resistance gene, *aphA-3,* or tetracycline (Tet) resistance gene, *tetM*, using gene splicing by overlap extension (SOEing) PCR (Ho et al., 1989) with the primers listed in Table S3. With the exception of Fig. 5, all Δ*ssaACB* experiments were completed with the Kan^R^ strain, JFP169. The plasmids pVA206 (Paik et al., 2005) and pJFP76 (Turner et al., 2009a) were the sources of the *aphA-3* and *tetM* genes, respectively. The Δ*relA*, Δ*relP,* and Δ*relQ* mutants were re-created for this study by amplifying the *aphA-3* gene and flanking DNA from the corresponding, previously created mutants (Xu et al., 2011) using primers from the same study. All PCR products were purified using a Qiagen MinElute PCR kit prior to transformation. Transformations employing antibiotic selection were performed using the protocol described previously (Paik et al., 2005). Briefly, an overnight culture of the parent strain was grown in BD Bacto™ Todd Hewitt broth with horse serum (Invitrogen), then diluted 200-fold and incubated at 37°C. Optical density (OD_600_) of tube cultures was determined using a ThermoScientific BioMate 3S UV-VIS spectrophotometer. Knockout construct DNA (100 ng) and *S. sanguinis* competence stimulating peptide (70 ng) were added to the culture (OD_600_ ~0.07) and incubated at 37°C for 1.5 h prior to selective plating on BHI (BD) agar plates with antibiotics. Kan (Sigma-Aldrich) or Tet (Sigma-Aldrich) was added to a final concentration of 500 and 5 μg mL^−1^, respectively. All plates were incubated for 24 h at 37°C under anaerobic conditions, where atmospheric composition was adjusted using a programmable Anoxomat™ Mark II jar-filling system (AIG, Inc.) and a palladium catalyst was included in the jars. The Δ*ssaACB*::*aphA-3* mutation was nonpolar, as confirmed by complementation (Murgas et al., 2020). All mutants were confirmed to have the expected composition by sequence analysis of the DNA flanking the insertion sites.

**Table 2.** Strains used in this study.

| Strain | Description | Source or Reference |
| --- | --- | --- |
| SK36 | Human oral plaque isolate | M. Killian, Aarhus University<br>Xu et al. (2007) |
| JFP132 | Δ*sodA::aphA-3* | Crump et al. (2014) |
| JFP169<sup>†</sup> | Δ*ssaACB::aphA-3* | This study |
| JFP173 | Δ*ssaACB::tetM* | This study |
| SSX_0250 | Δ*relA::aphA-3* | Xu et al. (2011) |
| JFP259 | Δ*relA::aphA-3* | This study |
| JFP260 | Δ*ssaACB::tetM* Δ*relA::aphA-3* | This study |
| SSX_1210 | $\Delta relQ::aphA-3$ | Xu et al. (2011) |
| JFP279 | $\Delta relQ::aphA-3$ | This study |
| JFP281 | $\Delta ssaACB::tetM \Delta relQ::aphA-3$ | This study |
| SSX_1795 | $\Delta relP::aphA-3$ | Xu et al. (2011) |
| JFP275 | $\Delta relP::aphA-3$ | This study |
| JFP277 | $\Delta ssaACB::tetM \Delta relP::aphA-3$ | This study |
| JFP276 | $\Delta relA::aphA-3 \Delta relP$ | This study |
| JFP278 | $\Delta ssaACB::tetM \Delta relA::aphA-3 \Delta relP$ | This study |
| JFP280 | $\Delta relA::aphA-3 \Delta relP \Delta relQ (rel^0)$ | This study |
| JFP282 | $\Delta ssaACB::tetM \Delta relA::aphA-3 \Delta relP \Delta relQ (rel^0)$ | This study |
| SK36 IFDC | *pheS\** *ermAM* | Cheng et al. (2018) |

<sup>†</sup>Designated $\Delta ssaACB$ throughout the manuscript

To generate the *rel*^0^ strain, Δ*relP* and Δ*relQ* mutations were made using a markerless mutation system described previously (Xie et al., 2011;Cheng et al., 2018). Briefly, the IFDC2 cassette was amplified from the *S. sanguinis* IFDC2 strain and combined with flanking region from *relP* using gene SOEing. The two parent Δ*relA* strains (WT and Δ*ssaACB*::*tetM* backgrounds) were then transformed as described above, plating on BHI agar plates containing 10 μg mL^−1^ erythromycin (Erm; Fisher Scientific). A gene SOEing product merging the two flanking regions of *relP* was then generated. This SOEing product was then used to transform the Erm^R^ colonies from the first transformation. Immediately prior to plating on agar plates containing 20 mM 4-chloro-phenylalanine (Sigma-Aldrich), the cells were washed twice with phosphate-buffered saline (PBS) to remove excess media. Resulting colonies were then screened for Erm sensitivity and sequenced to confirm removal of the desired gene and IFDC2 cassette. The process was then repeated for *relQ*, converting the two Δ*relA* Δ*relP* parent strains into *rel*^0^ strains.

Overnight BHI cultures (pre-cultures) were inoculated from single-use aliquots of cryopreserved cells by 1000-fold dilution. Antibiotics were included in mutant pre-cultures at the aforementioned concentrations. Cultures were then incubated for approximately 18 h at 6% O_2_ (Anoxomat jar set to 6% O_2_, 7% H_2_, 7% CO_2_ and 80% N_2_) at 37°C. To determine CFUs, samples were sonicated for 90 s using an ultrasonic homogenizer (Biologics, Inc) to disrupt chains prior to dilution in PBS and plated using an Eddy Jet 2 spiral plater (Neutec Group, Inc.). For static growth studies, tubes containing either 100% pooled rabbit serum (Gibco) or BHI with 100 μM EDTA (Invitrogen) were pre-incubated at 37°C in an atmosphere of 6% or 12% O_2_ (12% O_2_, 4.3% CO_2_, 4.3% H_2_). Each tube was then inoculated with a 10^−6^-fold dilution of the overnight pre-culture, as described above. For experiments depicted in Fig. S7B, 4 μM L-cysteine (Alfa Aesar) or reduced glutathione (ACROS Organics) was added to culture tubes immediately prior to inoculation. The inoculated tubes were returned to incubate at the same oxygen concentration. Cultures were removed after 24 h, sonicated, and diluted in PBS prior to plating on BHI agar. Plates were incubated for 24 h in 0% O_2_ prior to colony enumeration.

### 5.2 Fermentor growth conditions and sample collection

A BIOSTAT® B bioreactor (Sartorius Stedim) with a 1.5-L capacity UniVessel^®^ glass vessel was used for growth of 800-mL cultures at 37°C. Cultures were stirred at 250 rpm and pH was maintained by the automated addition of 2 N KOH (Fisher Chemical). A 40-mL overnight pre-culture of *S. sanguinis* was grown as described above and centrifuged for 10 minutes at 3,740 *x g* in an Allegra X-142 centrifuge at 4°C (Beckman-Coulter). The supernatant was discarded and the cells were resuspended in BHI prior to inoculation. The air flow was increased stepwise, based on the OD of the fermentor culture. At the peak OD, air flow was increased to 0.50 lpm, input flow of fresh BHI was set to 17% (~700 mL h^−1^) and output flow of waste was set to 34%. Cells were allowed to acclimate to the new conditions for 1 h. The T_−20_ sample was aseptically removed for total RNA isolation or ICP-OES analysis (described below). EDTA (Invitrogen) was introduced to the carboy 16 mins later (T_−4_) to achieve a final concentration of 100 μM. EDTA was then introduced directly to the vessel 4 mins later (T_0_), corresponding to the time at which EDTA from the carboy would reach the vessel, to achieve a final concentration of 100 μM. Samples were taken for each post-EDTA time point (T_10_, T_25_, T_50_). In some experiments, a divalent metal (Puratronic™ Alfa Aesar) was added to the carboy (T_66_) and vessel (T_70_) at a final concentration of 100 μM and samples were taken for ICP-OES at T_80_.

### 5.3 Total RNA isolation

For each RNA sample, 2 mL of fermentor culture was added to 4 mL RNAprotect Bacteria Reagent (Qiagen) and immediately vortexed for 10 sec. The sample was then incubated at room temperature for 5-90 min. The samples were then centrifuged for 10 min at 3,740 x *g* at 4°C. The supernatant was discarded and the samples stored at −80°C. RNA isolation and on-column DNase treatment were completed using the RNeasy Mini Kit and RNase-Free DNase Kit, respectively (Qiagen). RNA was eluted in 50 μL RNase-Free water (Qiagen). A second DNase treatment was then performed on the samples (Invitrogen). Total RNA was quantified and purity was assessed using a Nanodrop 2000 Spectrophotometer (ThermoScientific).

### 5.4 RNA-seq library preparation and sequencing

Total RNA quantity and integrity were determined using an Agilent Bioanalyzer RNA Pico assay. All samples passed quality control assessment with RNA Integrity Numbers (RIN) above 8. Two sequential rounds of ribosomal reduction were then performed on all samples using Illumina’s Ribo-Zero rRNA Removal Kit. The resulting depleted RNA was assessed using an Agilent Bioanalyzer RNA Pico assay to confirm efficient rRNA removal. Stranded RNA-seq library construction was then performed on the rRNA-depleted RNA using the Ultra II Directional RNA Library Prep Kit for Illumina (New England Biolabs), following manufacturer’s specifications for library construction and multiplexing. Final Illumina libraries were assessed for quality using an Agilent Bioanalyzer DNA High Sensitivity Assay and qPCR quantification was performed using NEBNext Library Quant kit for Illumina (New England Biolabs). Individual libraries were pooled equimolarly, and the final pool was sequenced on an Illumina MiSeq, with 2 x 150-bp paired-end reads. Demultiplexing was performed on the Illumina MiSeq’s on-board computer and resulting demultiplexed files uploaded to Illumina BaseSpace for data delivery. The University of Virginia Department of Biology Genomics Core Facility (Charlottesville, Virginia) completed all RNA-seq library preparation and sequencing.

### 5.5 RNA-seq analysis pipeline

Using Geneious 11.1 (https://www.geneious.com; RRID:SCR_010519), sequence reads were trimmed using the BBDuk Trimmer prior to mapping to a modified SK36 genome, in which the *ssaACB* operon was replaced with the *aphA-3* sequence. The locus tags are from the Genbank^®^ annotation (Benson et al., 2013) available at the time; the annotations were updated shortly before publication and the new locus tags are included in Table S1 for reference. PATRIC annotations (https://patricbrc.org/; RRID:SCR_004154) (Wattam et al., 2017) are also included for reference. Reads for each post-EDTA sample were compared to the corresponding pre-EDTA (T_−20_) sample using DESeq2 (Love et al., 2014) (RRID: SCR_015687) in Geneious to determine log_2_ fold changes and adjusted *P*-values. Principal component analysis was completed using R (version 3.6.1) and RStudio (version 1.2.5033-1) with Bioconductor (Bioconductor, RRID:SCR_006442) package pcaExplorer version 2.13.0 (Marini and Binder, 2019). Volcano plots were generated using R and RStudio with Bioconductor package EnhancedVolcano (Blighe et al., 2018). All DEGs were input into the DAVID database (https://david.ncifcrf.gov/summary.jsp; RRID:SCR_001881) (Dennis et al., 2003). The KEGG_pathway option was chosen for functional annotation clustering. The *P*-value shows the significance of pathway enrichment. Fig. 2C was generated using an R script (https://github.com/DrBinZhu/DAVID_FIG).

### 5.6 Inductively coupled plasma-optical emission spectroscopy

Additional 40-mL cell culture samples were collected from WT and Δ*ssaACB* cells at the same fermentor growth time points as described above. The cells were immediately centrifuged at 3,740 x *g* for 10 min at 4°C. The supernatant was decanted and the cell pellet was washed twice with cold cPBS (PBS treated with Chelex-100 resin (Bio-Rad) for 2 h, then filter sterilized and supplemented with EDTA to 1 mM). The pellet was then divided for subsequent acid digestion or protein concentration determination. Trace metal grade (TMG) nitric acid (15%) (Fisher Chemical) was added to one portion of the pellet. The pellet was digested using an Anton Paar microwave digestion system using a modified Organic B protocol: 120°C for 10 min, 180°C for 20 min, with the maximum temperature set to 180°C. The digested samples were then diluted 3-fold with Chelex-treated dH_2_O. Metal concentrations were determined using an Agilent 5110 inductively coupled plasma-optical emission spectrometer (ICP-OES). Concentrations were determined by comparison with a standard curve created with a 10 μg ml^−1^ multi-element standard (CMS-5; Inorganic Ventures) diluted in 5% TMG nitric acid. Pb (Inorganic Ventures) was used as an internal standard (10 μg ml^−1^). The other portion of the pellet was resuspended in PBS and mechanically lysed using a FastPrep-24 instrument with Lysing Matrix B tubes (MP Biomedicals) as described previously (Rhodes et al., 2014). Insoluble material was removed by centrifugation. Protein concentrations were determined using a bicinchoninic acid (BCA) Protein Assay Kit (Pierce) as recommended by the manufacturer, with bovine serum albumin as the standard. Absorbance was measured in a black, flat-bottom 96-well plate (Greiner) using a microplate reader (BioTek).

### 5.7 Hydrogen peroxide quantification

Culture supernatants without RNAprotect were collected at each time point and stored at −20°C. Hydrogen peroxide concentration was measured using a Fluorometric Hydrogen Peroxide Assay Kit (Sigma). Standards were prepared from 3% hydrogen peroxide provided with the kit as recommended by the manufacturer. Fluorescence was measured in a black-walled, flat-bottom 96-well plate (Greiner) using a microplate reader (BioTek).

### 5.8 Data analysis and presentation

All statistical tests, excluding RNA-seq DESeq2 calculations, were performed in GraphPad InStat (graphpad.com). Significance was determined by analysis of variance (ANOVA) as indicated in the figure legends. For ANOVA, a Tukey-Kramer test for multiple comparisons was used when *P* ≤ 0.05. DESeq2 calculations were completed in Geneious 11.1 or in the EnhancedVolcano R package, as indicated above. Confidence intervals (95%) of replicate samples were determined by the pcaExplorer R package. *P* values ≤ 0.05 were considered significant. All graphs and the heat map were constructed using GraphPad Prism (graphpad.com); Fig. 7 was made using Biorender.com.

## 7 Author Contributions

TP and TK designed the experiments and wrote all drafts of the manuscript. TP and KK performed the experiments. TP, KK, and BZ generated the figures. TP, TK, KK, BZ, and PX analyzed the data. All authors reviewed and approved the final version of the manuscript.

## 8 Contribution to the Field

The importance of manganese (Mn) for the survival and virulence of many bacterial pathogens, including streptococci, has been thoroughly documented. While several bacterial enzymes have been identified that require Mn as a cofactor, the full impact of Mn on the opportunistic endocarditis pathogen *Streptococcus sanguinis* has not been previously characterized. We employed a fermentor to maintain steady growth conditions, and to the best of our knowledge, the technique has not been used to examine metal-dependent gene expression in any streptococcal species. We observed that Mn depletion leads to changes in diverse systems with no apparent connection to the pathways of known Mn-cofactored enzymes. We also provide further evidence for a relationship between Mn and the regulation of carbon source utilization through carbon catabolite repression. The majority of the changes we observed have a potential connection to CcpA repression through putative carbon responsive elements identified upstream of the coding region. Here we confirm an earlier finding of glucose-independent catabolite repression, and for the first time, tie it to Mn depletion. This relationship will be investigated further in *S. sanguinis* and other related species to determine whether this is a species-specific phenotype.

## 10 Supplementary Material

Supplementary File: PDF

Supplementary Spreadsheets: Tables S1-S3

## 11 Acknowledgements

We gratefully acknowledge Dr. Shannon Green, Dr. Seon-Sook An, Brittany Spivey, and Rachel Korba for meaningful discussions and assistance with experiments. We appreciate Dr. Jody Turner (VCU Department of Chemistry) for advice and technical support regarding ICP-OES. We thank Drs. Jens Kreth and Nyssa Cullin (Oregon Health & Science University) for providing the IFDC2 *S. sanguinis* strain and corresponding protocol and Stephanie Neal for construction of JFP169.

## 12 Data Availability Statement

The data discussed in this publication have been deposited in NCBI's Gene Expression Omnibus (Edgar et al., 2002) and are accessible through GEO Series accession number GSE150593 (https://www.ncbi.nlm.nih.gov/geo/query/acc.cgi?acc=GSE150593).

## References

Ajdic, D., and Pham, V.T. (2007). Global transcriptional analysis of *Streptococcus mutans* sugar transporters using microarrays. J Bacteriol 189, 5049–5059.

Bai, Y., Shang, M., Xu, M., Wu, A., Sun, L., and Zheng, L. (2019). Transcriptome, phenotypic, and virulence analysis of *Streptococcus sanguinis* SK36 wild type and its CcpA-null derivative (D*ccpA*). Front Cell Infect Microbiol 9, 411.

Barnese, K., Gralla, E.B., Cabelli, D.E., and Valentine, J.S. (2008). Manganous phosphate acts as a superoxide dismutase. JACS 130, 4604–4606.

Barnese, K., Gralla, E.B., Valentine, J.S., and Cabelli, D.E. (2012). Biologically relevant mechanism for catalytic superoxide removal by simple manganese compounds. Proc Natl Acad Sci U S A 109, 6892–6897.

Bashore, T.M., Cabell, C., and Fowler, V., Jr. (2006). Update on infective endocarditis. Curr Probl Cardiol 31, 274–352.

Bayle, L., Chimalapati, S., Schoehn, G., Brown, J., Vernet, T., and Durmort, C. (2011). Zinc uptake by *Streptococcus pneumoniae* depends on both AdcA and AdcAII and is essential for normal bacterial morphology and virulence. Mol Microbiol 82, 904–916.

Belda-Ferre, P., Alcaraz, L.D., Cabrera-Rubio, R., Romero, H., Simon-Soro, A., Pignatelli, M., and Mira, A. (2012). The oral metagenome in health and disease. ISME J 6, 46–56.

Bensing, B.A., Li, L., Yakovenko, O., Wong, M., Barnard, K.N., Iverson, T.M., Lebrilla, C.B., Parrish, C.R., Thomas, W.E., Xiong, Y., and Sullam, P.M. (2019). Recognition of specific sialoglycan structures by oral streptococci impacts the severity of endocardial infection. PLoS Pathog 15, e1007896.

Bensing, B.A., Li, Q., Park, D., Lebrilla, C.B., and Sullam, P.M. (2018). Streptococcal Siglec-like adhesins recognize different subsets of human plasma glycoproteins: implications for infective endocarditis. Glycobiology 28, 601–611.

Benson, D.A., Cavanaugh, M., Clark, K., Karsch-Mizrachi, I., Lipman, D.J., Ostell, J., and Sayers, E.W. (2013). GenBank. Nucleic Acids Res 41, D36–42.

Bersch, B., Bougault, C., Roux, L., Favier, A., Vernet, T., and Durmort, C. (2013). New insights into histidine triad proteins: Solution structure of a *Streptococcus pneumoniae* PhtD domain and zinc transfer to AdcAII. PLoS One 8, e81168.

Bhubhanil, S., Chamsing, J., Sittipo, P., Chaoprasid, P., Sukchawalit, R., and Mongkolsuk, S. (2014). Roles of *Agrobacterium tumefaciens* membrane-bound ferritin (MbfA) in iron transport and resistance to iron under acidic conditions. Microbiology 160, 863–871.

Blighe, K., Rana, S., and Lewis, M. (2018). EnhancedVolcano: Publication-ready volcano plots with enhanced colouring and labeling [Online]. GitHub. Available: https://github.com/kevinblighe/EnhancedVolcano. [Accessed January 10, 2020].

Bor, D.H., Woolhandler, S., Nardin, R., Brusch, J., and Himmelstein, D.U. (2013). Infective endocarditis in the U.S., 1998-2009: a nationwide study. PLoS One 8, e60033.

Borovok, I., Gorovitz, B., Yanku, M., Schreiber, R., Gust, B., Chater, K., Aharonowitz, Y., and Cohen, G. (2004). Alternative oxygen-dependent and oxygen-independent ribonucleotide reductases in *Streptomyces*: cross-regulation and physiological role in response to oxygen limitation. Mol Microbiol 54, 1022–1035.

Brenot, A., King, K.Y., and Caparon, M.G. (2005). The PerR regulon in peroxide resistance and virulence of *Streptococcus pyogenes*. Mol Microbiol 55, 221–234.

Burne, R.A., and Chen, Y.-Y.M. (1998). The use of continuous flow bioreactors to explore gene expression and physiology of suspended and adherent populations of oral streptococci. Methods in Cell Science 20, 181–190.

Cahill, T.J., Baddour, L.M., Habib, G., Hoen, B., Salaun, E., Pettersson, G.B., Schafers, H.J., and Prendergast, B.D. (2017). Challenges in Infective Endocarditis. J Am Coll Cardiol 69, 325–344.

Chen, L., Ge, X., Dou, Y., Wang, X., Patel, J.R., and Xu, P. (2011). Identification of hydrogen peroxide production-related genes in *Streptococcus sanguinis* and their functional relationship with pyruvate oxidase. Microbiology 157, 13–20.

Chen, L., Ge, X., Wang, X., Patel, J.R., and Xu, P. (2012). SpxA1 involved in hydrogen peroxide production, stress tolerance and endocarditis virulence in *Streptococcus sanguinis*. PloS one 7, e40034–e40034.

Chen, Y.M., Chiang, Y.C., Tseng, T.Y., Wu, H.Y., Chen, Y.Y., Wu, C.H., and Chiu, C.H. (2019). Molecular and functional analysis of the type IV pilus gene cluster in *Streptococcus sanguinis* SK36. Appl Environ Microbiol 85.

Cheng, X., Redanz, S., Cullin, N., Zhou, X., Xu, X., Joshi, V., Koley, D., Merritt, J., and Kreth, J. (2018). Plasticity of the pyruvate node modulates hydrogen peroxide production and acid tolerance in multiple oral streptococci. Appl Environ Microbiol 84.

Colomer-Winter, C., Flores-Mireles, A.L., Kundra, S., Hultgren, S.J., and Lemos, J.A. (2019). (p)ppGpp and CodY promote *Enterococcus faecalis* virulence in a murine model of catheter-associated urinary tract infection. mSphere 4.

Colomer-Winter, C., Gaca, A.O., and Lemos, J.A. (2017). Association of metal homeostasis and (p)ppGpp regulation in the pathophysiology of *Enterococcus faecalis*. Infect Immun 85.

Cotruvo, J.A., Jr., and Stubbe, J. (2012). Metallation and mismetallation of iron and manganese proteins in vitro and in vivo: the class I ribonucleotide reductases as a case study. Metallomics 4, 1020–1036.

Couvin, D., Bernheim, A., Toffano-Nioche, C., Touchon, M., Michalik, J., Neron, B., Rocha, E.P.C., Vergnaud, G., Gautheret, D., and Pourcel, C. (2018). CRISPRCasFinder, an update of CRISRFinder, includes a portable version, enhanced performance and integrates search for Cas proteins. Nucleic Acids Res 46, W246–W251.

Crump, K.E., Bainbridge, B., Brusko, S., Turner, L.S., Ge, X., Stone, V., Xu, P., and Kitten, T. (2014). The relationship of the lipoprotein SsaB, manganese and superoxide dismutase in *Streptococcus sanguinis* virulence for endocarditis. Mol Microbiol 92, 1243–1259.

Das, S., Kanamoto, T., Ge, X., Xu, P., Unoki, T., Munro, C.L., and Kitten, T. (2009). Contribution of lipoproteins and lipoprotein processing to endocarditis virulence in *Streptococcus sanguinis*. J Bacteriol 191, 4166–4179.

Dayer, M., and Thornhill, M. (2018). Is antibiotic prophylaxis to prevent infective endocarditis worthwhile? J Infect Chemother 24, 18–24.

Dennis, G., Jr., Sherman, B.T., Hosack, D.A., Yang, J., Gao, W., Lane, H.C., and Lempicki, R.A. (2003). DAVID: Database for Annotation, Visualization, and Integrated Discovery. Genome Biol 4, P3.

Dintilhac, A., and Claverys, J.-P. (1997). The *adc* locus, which affects competence for genetic transformation in *Streptococcus pneumoniae*, encodes an ABC transporter with a putative lipoprotein homologous to a family of streptococcal adhesins. Res Microbiol 148, 119–131.

Dodds, D.R. (2017). Antibiotic resistance: A current epilogue. Biochem Pharmacol 134, 139–146.

Downey, J.S., Mashburn-Warren, L., Ayala, E.A., Senadheera, D.B., Hendrickson, W.K., Mccall, L.W., Sweet, J.G., Cvitkovitch, D.G., Spatafora, G.A., and Goodman, S.D. (2014). *In vitro* manganese-dependent cross-talk between *Streptococcus mutans* VicK and GcrR: implications for overlapping stress response pathways. PLoS One 9, e115975.

Edgar, R., Domrachev, M., and Lash, A.E. (2002). Gene Expression Omnibus: NCBI gene expression and hybridization array data repository. Nucleic Acids Research 30, 207–210.

Eijkelkamp, B.A., Mcdevitt, C.A., and Kitten, T. (2015). Manganese uptake and streptococcal virulence. Biometals 28, 491–508.

Eijkelkamp, B.A., Morey, J.R., Ween, M.P., Ong, C.L., Mcewan, A.G., Paton, J.C., and Mcdevitt, C.A. (2014). Extracellular zinc competitively inhibits manganese uptake and compromises oxidative stress management in *Streptococcus pneumoniae*. PLoS One 9, e89427.

Forner, L., Larsen, T., Kilian, M., and Holmstrup, P. (2006). Incidence of bacteremia after chewing, tooth brushing and scaling in individuals with periodontal inflammation. J Clin Periodontol 33, 401–407.

Fox, K.A., Ramesh, A., Stearns, J.E., Bourgogne, A., Reyes-Jara, A., Winkler, W.C., and Garsin, D.A. (2009). Multiple posttranscriptional regulatory mechanisms partner to control ethanolamine utilization in *Enterococcus faecalis*. Proc Natl Acad Sci U S A 106, 4435–4440.

Garcia-Mendoza, A., Liebana, J., Castillo, A.M., Higuera, A.D.L., and Piedrola, G. (1993). Evaluation of the capacity of oral streptococci to produce hydrogen peroxide. Journal of Medical Microbiology 39, 434–439.

Gaustad, P., and Havarstein, L.S. (1997). Competence pheromone in *Streptococcus sanguis*: Identification of the competence gene *comC* and the competence pheromone. Adv Exp Med Biol 418, 1019–1021.

Giacaman, R.A., Torres, S., Gomez, Y., Munoz-Sandoval, C., and Kreth, J. (2015). Correlation of Streptococcus mutans and Streptococcus sanguinis colonization and ex vivo hydrogen peroxide production in carious lesion-free and high caries adults. Arch Oral Biol 60, 154–159.

Gong, T., Zeng, J., Tang, B., Zhou, X., and Li, Y. (2020). CRISPR-Cas systems in oral microbiome: From immune defense to physiological regulation. Mol Oral Microbiol 35, 41–48.

Gorke, B., and Stulke, J. (2008). Carbon catabolite repression in bacteria: many ways to make the most out of nutrients. Nat Rev Microbiol 6, 613–624.

Gratani, F.L., Horvatek, P., Geiger, T., Borisova, M., Mayer, C., Grin, I., Wagner, S., Steinchen, W., Bange, G., Velic, A., Macek, B., and Wolz, C. (2018). Regulation of the opposing (p)ppGpp synthetase and hydrolase activities in a bifunctional RelA/SpoT homologue from *Staphylococcus aureus*. PLoS Genet 14, e1007514.

Griffen, A.L., Beall, C.J., Campbell, J.H., Firestone, N.D., Kumar, P.S., Yang, Z.K., Podar, M., and Leys, E.J. (2012). Distinct and complex bacterial profiles in human periodontitis and health revealed by 16S pyrosequencing. ISME J 6, 1176–1185.

Grinberg, I., Shteinberg, T., Gorovitz, B., Aharonowitz, Y., Cohen, G., and Borovok, I. (2006). The *Streptomyces* NrdR transcriptional regulator is a Zn ribbon/ATP cone protein that binds to the promoter regions of class Ia and class II ribonucleotide reductase operons. J Bacteriol 188, 7635–7644.

Groisman, E.A., Hollands, K., Kriner, M.A., Lee, E.J., Park, S.Y., and Pontes, M.H. (2013). Bacterial Mg^2+^ homeostasis, transport, and virulence. Annu Rev Genet 47, 625–646.

Gross, E.L., Beall, C.J., Kutsch, S.R., Firestone, N.D., Leys, E.J., and Griffen, A.L. (2012). Beyond *Streptococcus mutans*: dental caries onset linked to multiple species by 16S rRNA community analysis. PLoS One 7, e47722.

Gurung, I., Spielman, I., Davies, M.R., Lala, R., Gaustad, P., Biais, N., and Pelicic, V. (2016). Functional analysis of an unusual type IV pilus in the Gram-positive *Streptococcus sanguinis*. Mol Microbiol 99, 380–392.

Ho, S.N., Hunt, H.D., Horton, R.M., Pullen, J.K., and Pease, L.R. (1989). Site-directed mutagenesis by overlap extension using the polymerase chain reaction. Gene 77, 51–59.

Hogg, T., Mechold, U., Malke, H., Cashel, M., and Hilgenfeld, R. (2004). Conformational antagonism between opposing active sites in a bifunctional RelA/SpoT homolog modulates (p)ppGpp metabolism during the stringent response. Cell.

Irving, S.E., and Corrigan, R.M. (2018). Triggering the stringent response: signals responsible for activating (p)ppGpp synthesis in bacteria. Microbiology 164, 268–276.

Jakubovics, N.S., Smith, A.W., and Jenkinson, H.F. (2002). Oxidative stress tolerance is manganese (Mn^2+^)-regulated in *Streptococcus gordonii*. Microbiology 148, 3255–3263.

Jakubovics, N.S., and Valentine, R.A. (2009). A new direction for manganese homeostasis in bacteria: Identification of a novel efflux system in *Streptococcus pneumoniae*. Mol Microbiol 72, 1–4.

Jamil, M., Sultan, I., Gleason, T.G., Navid, F., Fallert, M.A., Suffoletto, M.S., and Kilic, A. (2019). Infective endocarditis: trends, surgical outcomes, and controversies. J Thorac Dis 11, 4875–4885.

Jeske, L., Placzek, S., Schomburg, I., Chang, A., and Schomburg, D. (2019). BRENDA in 2019: a European ELIXIR core data resource. Nucleic Acids Res 47, D542–D549.

Johnston, J.W., Briles, D.E., Myers, L.E., and Hollingshead, S.K. (2006). Mn^2+^-dependent regulation of multiple genes in *Streptococcus pneumoniae* through PsaR and the resultant impact on virulence. Infect Immun 74, 1171–1180.

Jourlin-Castelli, C., Mani, N., Nakano, M.M., and Sonenshein, A.L. (2000). CcpC, a novel regulator of the LysR family required for glucose repression of the *citB* gene in *Bacillus subtilis*. J Mol Biol 295, 865–878.

Kallio, A., Sepponen, K., Hermand, P., Denoel, P., Godfroid, F., and Melin, M. (2014). Role of pneumococcal histidine triad (Pht) proteins in attachment of *Streptococcus pneumoniae* to respiratory epithelial cells. Infect Immun 82, 1683–1691.

Kanehisa, M., and Goto, S. (2000). KEGG: Kyoto Encyclopedia of Genes and Genomes. Nucleic Acids Research 28, 27–30.

Kaspar, J., Kim, J.N., Ahn, S.J., and Burne, R.A. (2016). An essential role for (p)ppGpp in the integration of stress tolerance, peptide signaling, and competence development in *Streptococcus mutans*. Front Microbiol 7, 1162.

Kaval, K.G., and Garsin, D.A. (2018). Ethanolamine utilization in bacteria. mBio 9, e00066–00018.

Kaval, K.G., Gebbie, M., Goodson, J.R., Cruz, M.R., Winkler, W.C., and Garsina, D.A. (2019). Ethanolamine utilization and bacterial microcompartment formation are subject to carbon catabolite repression. J Bacteriol 201, e00703–00718.

Kazmierczak, K.M., Wayne, K.J., Rechtsteiner, A., and Winkler, M.E. (2009). Roles of *rel_Spn_* in stringent response, global regulation and virulence of serotype 2 *Streptococcus pneumoniae* D39. Mol Microbiol 72, 590–611.

Kehres, D.G., Lawyer, C.H., and Maguire, M.E. (1998). The CorA magnesium transporter gene family. Microbial & Comparative Genomics 3, 151–169.

Kehres, D.G., and Maguire, M.E. (2003). Emerging themes in manganese transport, biochemistry and pathogenesis in bacteria. FEMS Microbiol Rev 27, 263–290.

Kholy, K.E., Genco, R.J., and Van Dyke, T.E. (2015). Oral infections and cardiovascular disease. Trends Endocrinol Metab 26, 315–321.

Kim, J.N., Ahn, S.J., Seaton, K., Garrett, S., and Burne, R.A. (2012). Transcriptional organization and physiological contributions of the *relQ* operon of *Streptococcus mutans*. J Bacteriol 194, 1968–1978.

Kim, S.A., Punshon, T., Lanzirotti, A., Li, L., Alonso, J.M., Ecker, J.R., Kaplan, J., and Guerinot, M.L. (2006). Localization of iron in *Arabidopsis* seed requires the vacuolar membrane transporter VIT1. Science 314, 1295–1298.

Kinane, D.F., Riggio, M.P., Walker, K.F., Mackenzie, D., and Shearer, B. (2005). Bacteraemia following periodontal procedures. J Clin Periodontol 32, 708–713.

Kolenbrander, P.E., Palmer, R.J., Jr., Periasamy, S., and Jakubovics, N.S. (2010). Oral multispecies biofilm development and the key role of cell-cell distance. Nat Rev Microbiol 8, 471–480.

Kreth, J., Merritt, J., Shi, W., and Qi, F. (2005). Competition and coexistence between *Streptococcus mutans* and *Streptococcus sanguinis* in the dental biofilm. J Bacteriol 187, 7193–7203.

Kreth, J., Zhang, Y., and Herzberg, M.C. (2008). Streptococcal antagonism in oral biofilms: *Streptococcus sanguinis* and *Streptococcus gordonii* interference with *Streptococcus mutans*. J Bacteriol 190, 4632–4640.

Kuipers, K., Gallay, C., Martinek, V., Rohde, M., Martinkova, M., Van Der Beek, S.L., Jong, W.S., Venselaar, H., Zomer, A., Bootsma, H., Veening, J.W., and De Jonge, M.I. (2016). Highly conserved nucleotide phosphatase essential for membrane lipid homeostasis in *Streptococcus pneumoniae*. Mol Microbiol 101, 12–26.

Labarbuta, P., Duckett, K., Botting, C.H., Chahrour, O., Malone, J., Dalton, J.P., and Law, C.J. (2017). Recombinant vacuolar iron transporter family homologue PfVIT from human malaria-causing *Plasmodium falciparum* is a Fe^2+^/H^+^ exchanger. Sci Rep 7, 1–10.

Lee, S., Kim, K.-K., and Choe, S.-J. (2001). Binding of oral streptococci to human fibrinogen. Oral Microbiol Immunol 16, 88–93.

Lemos, J.A., Lin, V.K., Nascimento, M.M., Abranches, J., and Burne, R.A. (2007). Three gene products govern (p)ppGpp production by *Streptococcus mutans*. Mol Microbiol 65, 1568–1581.

Li, L., Chen, O.S., Mcvey Ward, D., and Kaplan, J. (2001). CCC1 is a transporter that mediates vacuolar iron storage in yeast. J Biol Chem 276, 29515–29519.

Li, P., Gu, Q., Wang, Y., Yu, Y., Yang, L., and Chen, J.V. (2017). Novel vitamin B_12_-producing *Enterococcus* spp. and preliminary *in vitro* evaluation of probiotic potentials. Appl Microbiol Biotechnol 101, 6155–6164.

Liochev, S.I., and Fridovich, I. (2004). Carbon dioxide mediates Mn(II)-catalyzed decomposition of hydrogen peroxide and peroxidation reactions. Proc Natl Acad Sci U S A 101, 12485–12490.

Lisher, J.P., Higgins, K.A., Maroney, M.J., and Giedroc, D.P. (2013). Physical characterization of the manganese-sensing pneumococcal surface antigen repressor from *Streptococcus pneumoniae*. Biochemistry 52, 7689–7701.

Lockhart, P.B., Brennan, M.T., Sasser, H.C., Fox, P.C., Paster, B.J., and Bahrani-Mougeot, F.K. (2008). Bacteremia associated with toothbrushing and dental extraction. Circulation 117, 3118–3125.

Love, M.I., Huber, W., and Anders, S. (2014). Moderated estimation of fold change and dispersion for RNA-seq data with DESeq2. Genome Biol 15, 550.

Makarova, K.S., Wolf, Y.I., Alkhnbashi, O.S., Costa, F., Shah, S.A., Saunders, S.J., Barrangou, R., Brouns, S.J., Charpentier, E., Haft, D.H., Horvath, P., Moineau, S., Mojica, F.J., Terns, R.M., Terns, M.P., White, M.F., Yakunin, A.F., Garrett, R.A., Van Der Oost, J., Backofen, R., and Koonin, E.V. (2015). An updated evolutionary classification of CRISPR-Cas systems. Nat Rev Microbiol 13, 722–736.

Makhlynets, O., Boal, A.K., Rhodes, D.V., Kitten, T., Rosenzweig, A.C., and Stubbe, J. (2014). *Streptococcus sanguinis* class Ib ribonucleotide reductase: high activity with both iron and manganese cofactors and structural insights. J Biol Chem 289, 6259–6272.

Manzoor, I., Shafeeq, S., Afzal, M., and Kuipers, O.P. (2015). The regulation of the AdcR regulon in *Streptococcus pneumoniae* depends both on Zn^2+^- and Ni^2+^-availability. Front Cell Infect Microbiol 5, 1–11.

Marini, F., and Binder, H. (2019). pcaExplorer: an R/Bioconductor package for interacting with RNA-seq principal components. BMC Bioinformatics 20, 1–8.

Martin, J.E., and Giedroc, D.P. (2016). Functional determinants of metal ion transport and selectivity in paralogous cation diffusion facilitator transporters CzcD and MntE in *Streptococcus pneumoniae*. J Bacteriol 198, 1066–1076.

Martin, J.E., Le, M.T., Bhattarai, N., Capdevila, D.A., Shen, J., Winkler, M.E., and Giedroc, D.P. (2019). A Mn-sensing riboswitch activates expression of a Mn2+/Ca2+ ATPase transporter in *Streptococcus*. Nucleic Acids Res 47, 6885–6899.

Martin, J.E., Lisher, J.P., Winkler, M.E., and Giedroc, D.P. (2017). Perturbation of manganese metabolism disrupts cell division in *Streptococcus pneumoniae*. Mol Microbiol 104, 334–348.

Mechold, U., Gentry, D., Cashel, M., Steiner, K., and Malke, H. (1996). Functional analysis of a *relA/spoT* gene homolog from *Streptococcus equisimilis*. Journal of Bacteriology 178, 1401–1411.

Moore, C.M., Gaballa, A., Hui, M., Ye, R.W., and Helmann, J.D. (2005). Genetic and physiological responses of *Bacillus subtilis* to metal ion stress. Mol Microbiol 57, 27–40.

Moraes, J.J., Stipp, R.N., Harth-Chu, E.N., Camargo, T.M., Hofling, J.F., and Mattos-Graner, R.O. (2014). Two-component system VicRK regulates functions associated with establishment of *Streptococcus sanguinis* in biofilms. Infect Immun 82, 4941–4951.

Moreillon, P., and Que, Y.-A. (2004). Infective endocarditis. The Lancet 363, 139–149.

Moreillon, P., Que, Y.A., and Bayer, A.S. (2002). Pathogenesis of streptococcal and staphylococcal endocarditis. Infect Dis Clin North Am 16, 297–318.

Morona, J.K., Morona, R., Miller, D.C., and Paton, J.C. (2002). *Streptococcus pneumoniae* capsule biosynthesis protein CpsB is a novel manganese-dependent phosphotyrosine-protein phosphatase. J Bacteriol 184, 577–583.

Murgas, C.J., Green, S.P., Forney, A.K., Korba, R.M., An, S.S., Kitten, T., and Lucas, H.R. (2020). Intracellular metal speciation in *Streptococcus sanguinis* establishes SsaACB as critical for redox maintenance. ACS Infect Dis.

Nascimento, M.M., Lemos, J.A., Abranches, J., Lin, V.K., and Burne, R.A. (2008). Role of RelA of *Streptococcus mutans* in global control of gene expression. J Bacteriol 190, 28–36.

Nies, D.H. (1992). CzcR and CzcD, gene products affecting regulation of resistance to cobalt, zinc, and cadmium (*czc* system) in *Alcaligenes eutrophus*. Journal of Bacteriology 174, 8102–8110.

Novichkov, P.S., Kazakov, A.E., Ravcheev, D.A., Leyn, S.A., Kovaleva, G.Y., Sutormin, R.A., Kazanov, M.D., Riehl, W., Arkin, A.P., Dubchak, I., and Rodionov, D.A. (2013). RegPrecise 3.0 - A resource for genome-scale exploration of transcriptional regulation in bacteria. BMC Genomics 14, 745.

Ogunniyi, A.D., Mahdi, L.K., Jennings, M.P., Mcewan, A.G., Mcdevitt, C.A., Van Der Hoek, M.B., Bagley, C.J., Hoffmann, P., Gould, K.A., and Paton, J.C. (2010). Central role of manganese in regulation of stress responses, physiology, and metabolism in *Streptococcus pneumoniae*. J Bacteriol 192, 4489–4497.

Paik, S., Senty, L., Das, S., Noe, J.C., Munro, C.L., and Kitten, T. (2005). Identification of virulence determinants for endocarditis in *Streptococcus sanguinis* by signature-tagged mutagenesis. Infect Immun 73, 6064–6074.

Panosian, T.D., Nannemann, D.P., Watkins, G.R., Phelan, V.V., Mcdonald, W.H., Wadzinski, B.E., Bachmann, B.O., and Iverson, T.M. (2011). *Bacillus cereus* phosphopentomutase is an alkaline phosphatase family member that exhibits an altered entry point into the catalytic cycle. J Biol Chem 286, 8043–8054.

Papp-Wallace, K.M., and Maguire, M.E. (2006). Manganese transport and the role of manganese in virulence. Annu Rev Microbiol 60, 187–209.

Parker, M.W., and Blake, C.C.F. (1988). Iron- and manganese-containing superoxide dismutases can be distinguished by analysis of their primary structures. FEBS Lett 229, 377–382.

Parraga Solorzano, P.K., Yao, J., Rock, C.O., and Kehl-Fie, T.E. (2019). Disruption of glycolysis by nutritional immunity activates a two-component system that coordinates a metabolic and antihost response by *Staphylococcus aureus*. mBio 10.

Perrin, D.D., and Dempsey, B. (1974). Buffers for pH and Metal Ion Control. Bristol: J. W. Arrowsmith Ltd.

Plumptre, C.D., Eijkelkamp, B.A., Morey, J.R., Behr, F., Counago, R.M., Ogunniyi, A.D., Kobe, B., O'mara, M.L., Paton, J.C., and Mcdevitt, C.A. (2014). AdcA and AdcAII employ distinct zinc acquisition mechanisms and contribute additively to zinc homeostasis in *Streptococcus pneumoniae*. Mol Microbiol 91, 834–851.

Potter, A.J., Trappetti, C., and Paton, J.C. (2012). *Streptococcus pneumoniae* uses glutathione to defend against oxidative stress and metal ion toxicity. J Bacteriol 194, 6248–6254.

Poyart, C., Quesne, G., Coulon, S., Berche, P., and Trieu-Cuot, P. (1988). Identification of streptococci to species level by sequencing the gene encoding the manganese-dependent superoxide dismutase. Jour Clin Microbiol 36, 41–47.

Pulliainen, A.T., Haataja, S., Kahkonen, S., and Finne, J. (2003). Molecular basis of H_2_O_2_ resistance mediated by streptococcal Dpr: Demonstration of the functional involvement of the putative ferroxidase center by site-directed mutagenesis in *Streptococcus suis*. J Biol Chem 278, 7996–8005.

Quan, T.P., Muller-Pebody, B., Fawcett, N., Young, B.C., Minaji, M., Sandoe, J., Hopkins, S., Crook, D., Peto, T., Johnson, A.P., and Walker, A.S. (2020). Investigation of the impact of the NICE guidelines regarding antibiotic prophylaxis during invasive dental procedures on the incidence of infective endocarditis in England: an electronic health records study. BMC Med 18, 84.

Radin, J.N., Kelliher, J.L., Parraga Solorzano, P.K., and Kehl-Fie, T.E. (2016). The two-component system ArlRS and alterations in metabolism enable *Staphylococcus aureus* to resist calprotectin-induced manganese starvation. PLoS Pathog 12, e1006040.

Redanz, S., Masilamani, R., Cullin, N., Giacaman, R.A., Merritt, J., and Kreth, J. (2018a). Distinct regulatory role of carbon catabolite protein A (CcpA) in oral streptococcal *spxB* expression. J Bacteriol 200.

Redanz, S., Masilamani, R., Cullin, N., Giacaman, R.A., Merritt, J., and Kreth, J. (2018b). Distinct regulatory role of carbon catabolite protein A (CcpA) in oral streptococcal *spxB* expression. Journal of Bacteriology 200, e00619–00617.

Rhodes, D.V., Crump, K.E., Makhlynets, O., Snyder, M., Ge, X., Xu, P., Stubbe, J., and Kitten, T. (2014). Genetic characterization and role in virulence of the ribonucleotide reductases of *Streptococcus sanguinis*. J Biol Chem 289, 6273–6287.

Rodionov, D.A., and Gelfand, M.S. (2005). Identification of a bacterial regulatory system for ribonucleotide reductases by phylogenetic profiling. Trends Genet 21, 385–389.

Rodriguez, A.M., Callahan, J.E., Fawcett, P., Ge, X., Xu, P., and Kitten, T. (2011). Physiological and molecular characterization of genetic competence in *Streptococcus sanguinis*. Mol Oral Microbiol 26, 99–116.

Ronneau, S., and Hallez, R. (2019). Make and break the alarmone: regulation of (p)ppGpp synthetase/hydrolase enzymes in bacteria. FEMS Microbiol Rev 43, 389–400.

Rosch, J.W., Gao, G., Ridout, G., Wang, Y.D., and Tuomanen, E.I. (2009). Role of the manganese efflux system *mntE* for signalling and pathogenesis in *Streptococcus pneumoniae*. Mol Microbiol 72, 12–25.

Seaton, K., Ahn, S.J., Sagstetter, A.M., and Burne, R.A. (2011). A transcriptional regulator and ABC transporters link stress tolerance, (p)ppGpp, and genetic competence in *Streptococcus mutans*. J Bacteriol 193, 862–874.

Serbanescu, M.A., Cordova, M., Krastel, K., Flick, R., Beloglazova, N., Latos, A., Yakunin, A.F., Senadheera, D.B., and Cvitkovitch, D.G. (2015). Role of the *Streptococcus mutans* CRISPR-Cas systems in immunity and cell physiology. J Bacteriol 197, 749–761.

Shafeeq, S., Kloosterman, T.G., and Kuipers, O.P. (2011). Transcriptional response of *Streptococcus pneumoniae* to Zn^2+^ limitation and the repressor/activator function of AdcR. Metallomics 3, 609–618.

Sheppard, D., Berry, J.L., Denise, R., Rocha, E.P.C., Matthews, S., and Pelicic, V. (2020). The major subunit of widespread competence pili exhibits a novel and conserved type IV pilin fold. J Biol Chem.

Shields, R.C., Walker, A.R., Maricic, N., Chakraborty, B., Underhill, S.a.M., and Burne, R.A. (2020). Repurposing the *Streptococcus mutans* CRISPR-Cas9 system to understand essential gene function. PLoS Pathog 16, e1008344.

Silver, J.G., Martin, A.W., and Mcbride, B.C. (1977). Experimental transient bacteraemias in human subjects with varying degrees of plaque accumulation and gingival inflammation. J Clin Periodontol 4, 92–99.

Silver, J.G., Martin, A.W., and Mcbride, B.C. (1979). Experimental transient bacteraemias in human subjects with clinically healthy gingivae. J Clin Periodontol 6, 33–36.

Skov Sørensen, U.B., Yao, K., Yang, Y., Tettelin, H., and Kilian, M. (2016). Capsular polysaccharide expression in commensal *Streptococcus* species: Genetic and antigenic similarities to *Streptococcus pneumoniae*. mBio 7, e01844–01816.

Socransky, S.S., Manganiello, A.D., Propas, D., Oram, V., and Houte, J.V. (1977). Bacteriological studies of developing supragingival dental plaque. Journal Periodontal Research 12, 90–106.

Spellerberg, B., Cundell, D.R., Sandros, J., Pearce, B.J., Idanpaan-Heikkila, I., Rosenow, C., and Masure, H.R. (1996). Pyruvate oxidase, as a determinant of virulence in *Streptococcus pneumoniae*. Mol Microbiol 19, 803–813.

Sreenivasan, P.K., Tischio-Bereski, D., and Fine, D.H. (2017). Reduction in bacteremia after brushing with a triclosan/copolymer dentifrice-A randomized clinical study. J Clin Periodontol 44, 1020–1028.

Stingu, C.S., Eschrich, K., Rodloff, A.C., Schaumann, R., and Jentsch, H. (2008). Periodontitis is associated with a loss of colonization by *Streptococcus sanguinis*. J Med Microbiol 57, 495–499.

Thornhill, M.H., Gibson, T.B., Cutler, E., Dayer, M.J., Chu, V.H., Lockhart, P.B., O'gara, P.T., and Baddour, L.M. (2018). Antibiotic prophylaxis and incidence of endocarditis before and after the 2007 AHA recommendations. J Am Coll Cardiol 72, 2443–2454.

Torrents, E., Grinberg, I., Gorovitz-Harris, B., Lundstrom, H., Borovok, I., Aharonowitz, Y., Sjoberg, B.M., and Cohen, G. (2007). NrdR controls differential expression of the *Escherichia coli* ribonucleotide reductase genes. J Bacteriol 189, 5012–5021.

Tozzi, M.G., Camici, M., Mascia, L., Sgarrella, F., and Ipata, P.L. (2006). Pentose phosphates in nucleoside interconversion and catabolism. FEBS J 273, 1089–1101.

Tsunashima, H., Miyake, K., Motono, M., and Iijima, S. (2012). Organization of the capsule biosynthesis gene locus of the oral streptococcus *Streptococcus anginosus*. J Biosci Bioeng 113, 271–278.

Turner, L.S., Das, S., Kanamoto, T., Munro, C.L., and Kitten, T. (2009a). Development of genetic tools for *in vivo* virulence analysis of *Streptococcus sanguinis*. Microbiology 155, 2573–2582.

Turner, L.S., Kanamoto, T., Unoki, T., Munro, C.L., Wu, H., and Kitten, T. (2009b). Comprehensive evaluation of *Streptococcus sanguinis* cell wall-anchored proteins in early infective endocarditis. Infect Immun 77, 4966–4975.

Warner, J.B., and Lolkema, J.S. (2003). CcpA-dependent carbon catabolite repression in bacteria. Microbiology and Molecular Biology Reviews 67, 475–490.

Wattam, A.R., Davis, J.J., Assaf, R., Boisvert, S., Brettin, T., Bun, C., Conrad, N., Dietrich, E.M., Disz, T., Gabbard, J.L., Gerdes, S., Henry, C.S., Kenyon, R.W., Machi, D., Mao, C., Nordberg, E.K., Olsen, G.J., Murphy-Olson, D.E., Olson, R., Overbeek, R., Parrello, B., Pusch, G.D., Shukla, M., Vonstein, V., Warren, A., Xia, F., Yoo, H., and Stevens, R.L. (2017). Improvements to PATRIC, the all-bacterial bioinformatics database and analysis resource center. Nucleic Acids Res 45, D535–D542.

Widmer, E., Que, Y.-A., Entenza, J.M., and Moreillon, P. (2006). New concepts in the pathophysiology of infective endocarditis. Current Infectious Disease Reports 8, 271–279.

Wilson, W., Taubert, K.A., Gewitz, M., Lockhart, P.B., Baddour, L.M., Levison, M., Bolger, A., Cabell, C.H., Takahashi, M., Baltimore, R.S., Newburger, J.W., Strom, B.L., Tani, L.Y., Gerber, M., Bonow, R.O., Pallasch, T., Shulman, S.T., Rowley, A.H., Burns, J.C., Ferrieri, P., Gardner, T., Goff, D., and Durack, D.T. (2007). Prevention of infective endocarditis: Guidelines from the American Heart Association. Circulation 116, 1736–1754.

Xie, Z., Okinaga, T., Qi, F., Zhang, Z., and Merritt, J. (2011). Cloning-independent and counterselectable markerless mutagenesis system in *Streptococcus mutans*. Appl Environ Microbiol 77, 8025–8033.

Xu, P., Alves, J.M., Kitten, T., Brown, A., Chen, Z., Ozaki, L.S., Manque, P., Ge, X., Serrano, M.G., Puiu, D., Hendricks, S., Wang, Y., Chaplin, M.D., Akan, D., Paik, S., Peterson, D.L., Macrina, F.L., and Buck, G.A. (2007). Genome of the opportunistic pathogen *Streptococcus sanguinis*. J Bacteriol 189, 3166–3175.

Xu, P., Ge, X., Chen, L., Wang, X., Dou, Y., Xu, J.Z., Patel, J.R., Stone, V., Trinh, M., Evans, K., Kitten, T., Bonchev, D., and Buck, G.A. (2011). Genome-wide essential gene identification in *Streptococcus sanguinis*. Sci Rep 1, 1–9.

Xu, Y., Itzek, A., and Kreth, J. (2014). Comparison of genes required for H_2_O_2_ resistance in *Streptococcus gordonii* and *Streptococcus sanguinis*. Microbiology 160, 2627–2638.

Yamamoto, Y., Higuchi, M., Poole, L.B., and Kamio, Y. (2000). Role of the *dpr* product in oxygen tolerance in *Streptococcus mutans*. J Bacteriol 182, 3740–3747.

Yamamoto, Y., Poole, L.B., Hantgan, R.R., and Kamio, Y. (2002). An iron-binding protein, Dpr, from *Streptococcus mutans* prevents iron-dependent hydroxyl radical formation *in vitro*. J Bacteriol 184, 2931–2939.

Yang, N., Xie, S., Tang, N.Y., Choi, M.Y., Wang, Y., and Watt, R.M. (2019). The Ps and Qs of alarmone synthesis in *Staphylococcus aureus*. PLoS One 14, e0213630.

Zeng, L., Chen, L., and Burne, R.A. (2018). Preferred hexoses influence long-term memory in and induction of lactose catabolism by *Streptococcus mutans*. Applied and Environmental Microbiology 84, e00864–00818.

Zhang, Y.M., Liu, J.K., Shouri, M.R., and Wong, T.Y. (2006). Characterization of a Mn-dependent fructose-1,6-bisphosphate aldolase in *Deinococcus radiodurans*. Biometals 19, 31–37.

Zheng, L., Chen, Z., Itzek, A., Ashby, M., and Kreth, J. (2011). Catabolite control protein A controls hydrogen peroxide production and cell death in *Streptococcus sanguinis*. J Bacteriol 193, 516–526.

