## Supplemental File 1 for "Manganese depletion leads to multisystem changes in the transcriptome of the opportunistic pathogen *Streptococcus sanguinis*"

### *Supplementary Material*

#### Supplementary Figures

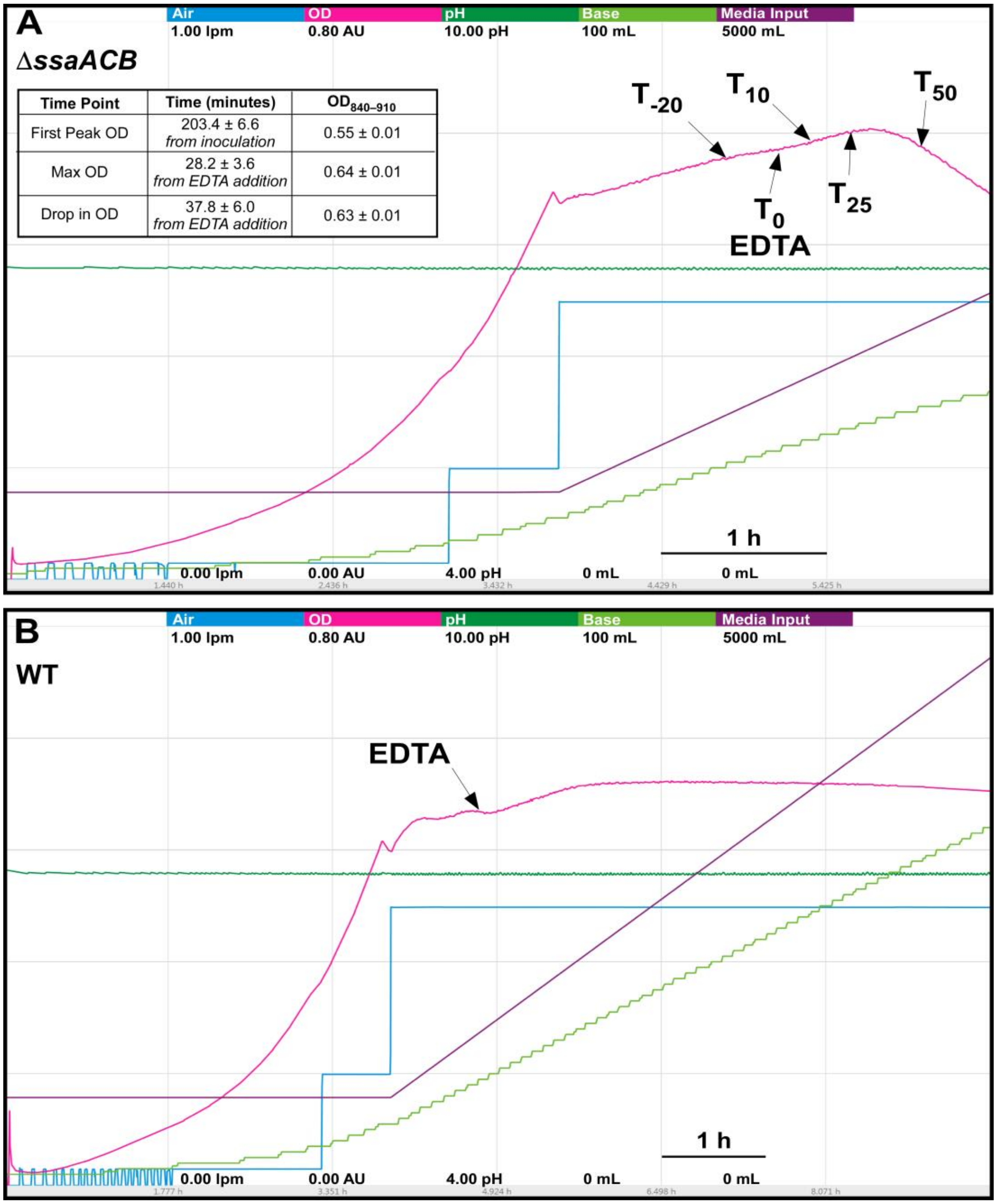


**Figure S1. Aerobic fermentor growth of Δ*ssaACB* and WT strains.**
Representative charts for fermentor growth of *S. sanguinis* (**A**) Δ*ssaACB* and (**B**) WT cells. Each color represents a different parameter: cyan - air flow (liters per min; lpm), pink - optical density (840-910 nm; absorbance units; AU), dark green - pH, light green - base input (KOH), purple - media input (total volume). Each color represents a different parameter as labeled at the top of the figure. The scale for each parameter is indicated by the values under each respective parameter label (minimum at the bottom, maximum at the top). The time scale is indicated by the bar in the bottom right portion of each chart. Cells were grown under aerobic conditions with EDTA added 80 mins ($T_0$) after the media input and output pumps were turned on and the air flow was set to 0.5 lpm. Each sample time point is labeled.

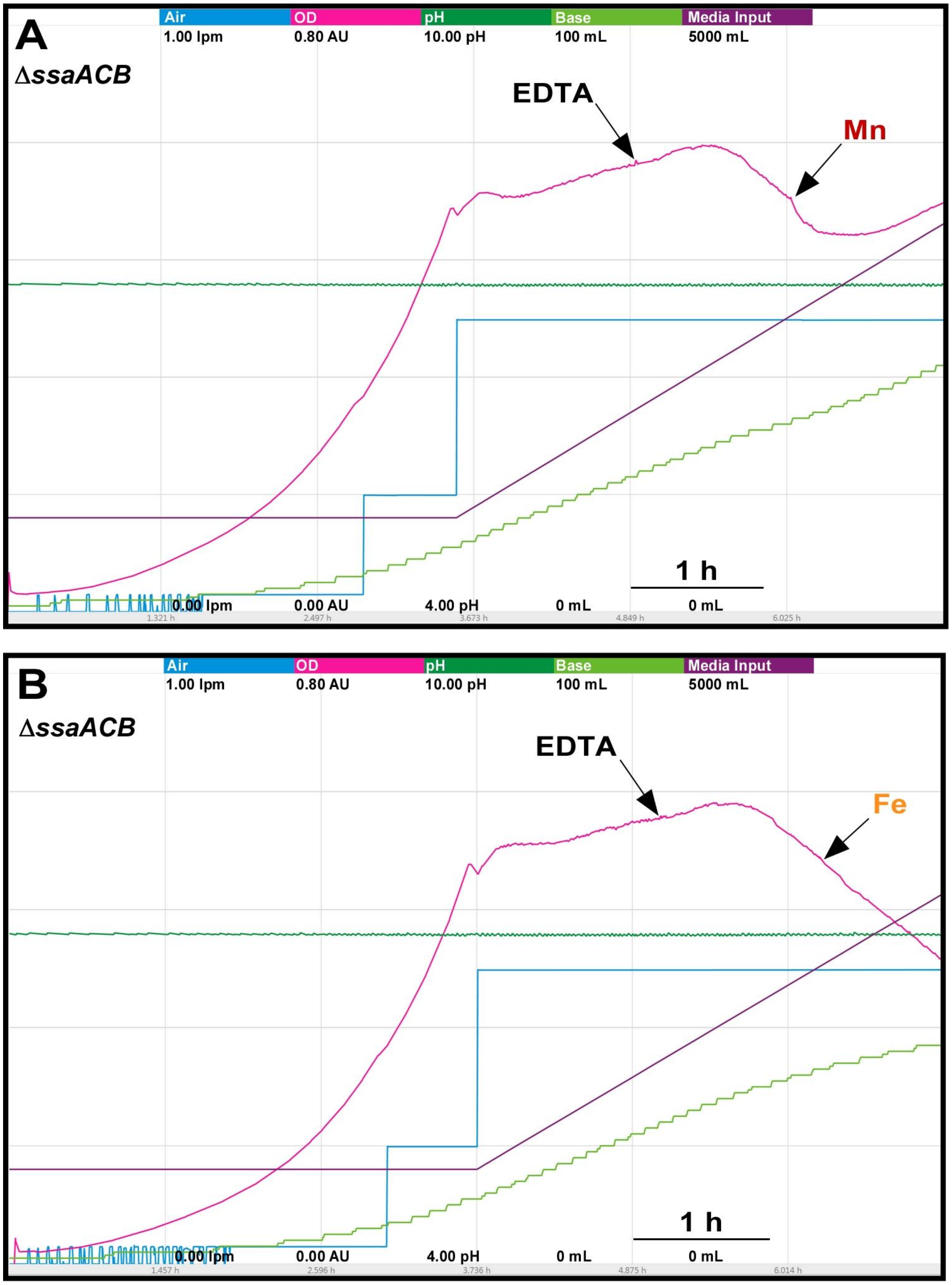


**Figure S2. Addition of metals to fermentor-grown Δ*ssaACB* cells post-EDTA.**
Fermentor growth of Δ*ssaACB* with the addition of 100 µM EDTA at $T_0$ as described previously, with 100 µM of either (**A**) Mn(II)$SO_4$ or (**B)** Fe(II)$SO_4$ added at $T_{70}$. Colors and labels are as in Fig. S1. The time scale is indicated by the bar in the bottom right portion of the each chart. Each chart is representative of at least three replicates.

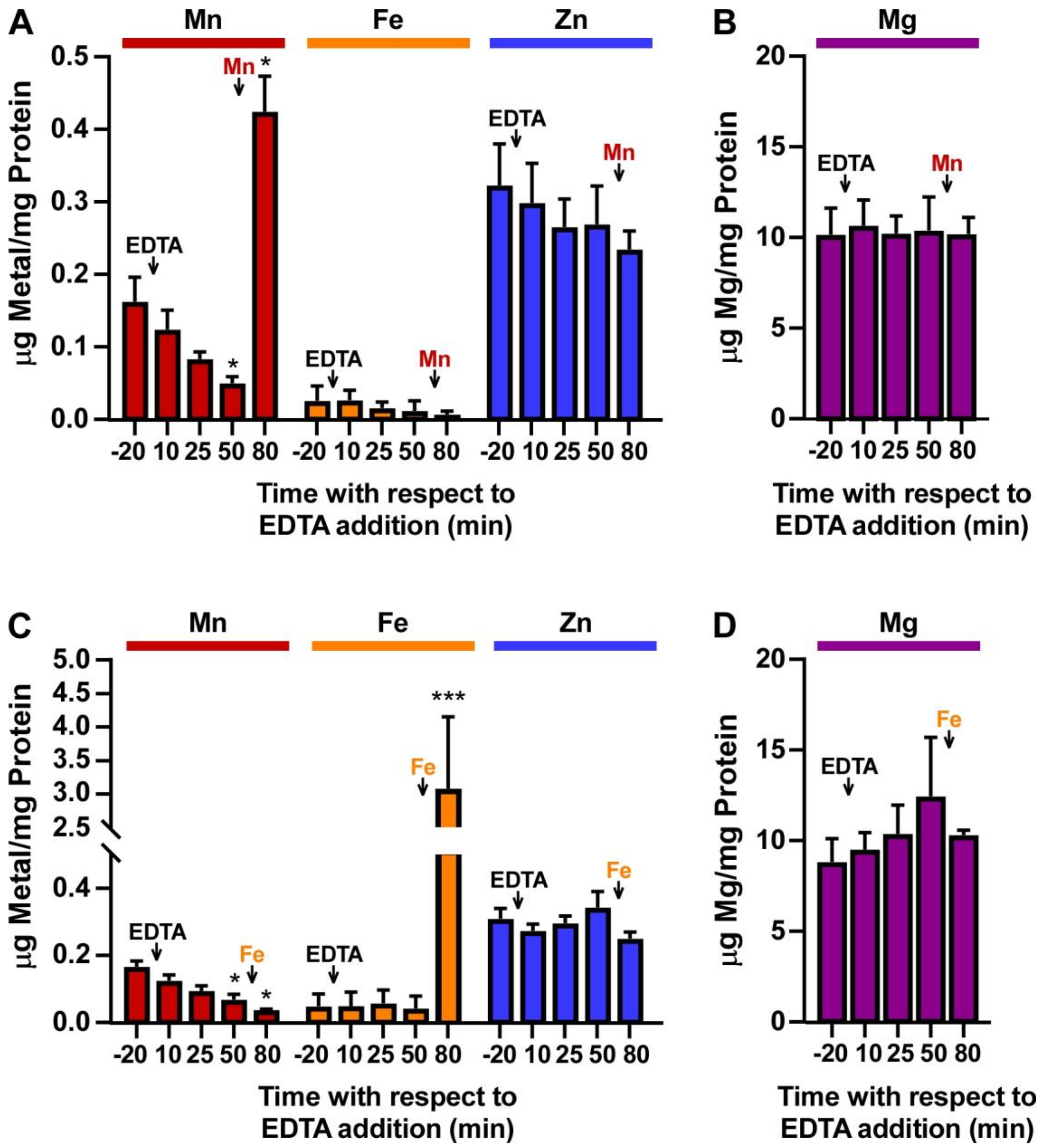


**Figure S3. Metal content of fermentor-grown Δ*ssaACB* cells post-EDTA and metal supplementation.** Samples of Δ*ssaACB* cells were collected from the fermentor at each time point and analyzed for cellular metal content using ICP-OES. The $T_{80}$ time point is 10 mins after the addition of 100 µM of either (**A-B**) Mn(II)$SO_4$ or (**C-D)** Fe(II)$SO_4$ added at $T_{70}$ as depicted in Figure S2. Means and standard deviations of three replicates are shown. Significance was determined for each metal by repeated measures ANOVA or one-way ANOVA if matching was not effective. A Tukey-Kramer multiple comparisons test was used for comparison to $T_{-20}$ for each metal; *$P \leq 0.05$, ***$P \leq 0.0001$.

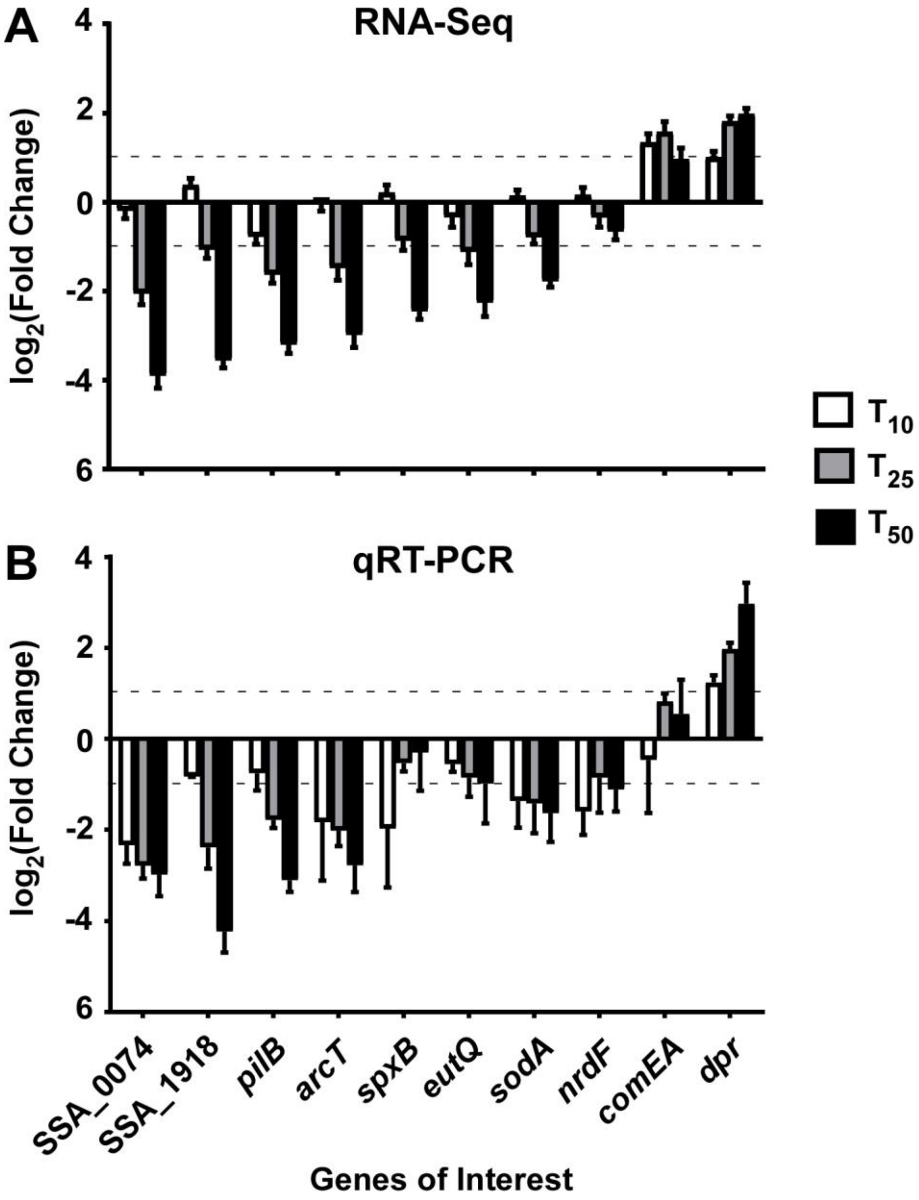


**Figure S4. Validation of RNA-Seq trends using qRT-PCR.**

(**A**) The average $\log_2$ fold change values of select genes of interest from the DESeq2 RNA-Seq analysis, comparing each post-EDTA time point to $T_{-20}$. The average is from four biological replicates. (**B**) $\text{Log}_2$ fold change values of the same genes as determined by qRT-PCR of two additional Δ*ssaACB* fermentor run samples. Mean and standard error for each time point are depicted. Horizontal dashed lines indicate $\log_2$ fold changes in expression of ± 1.

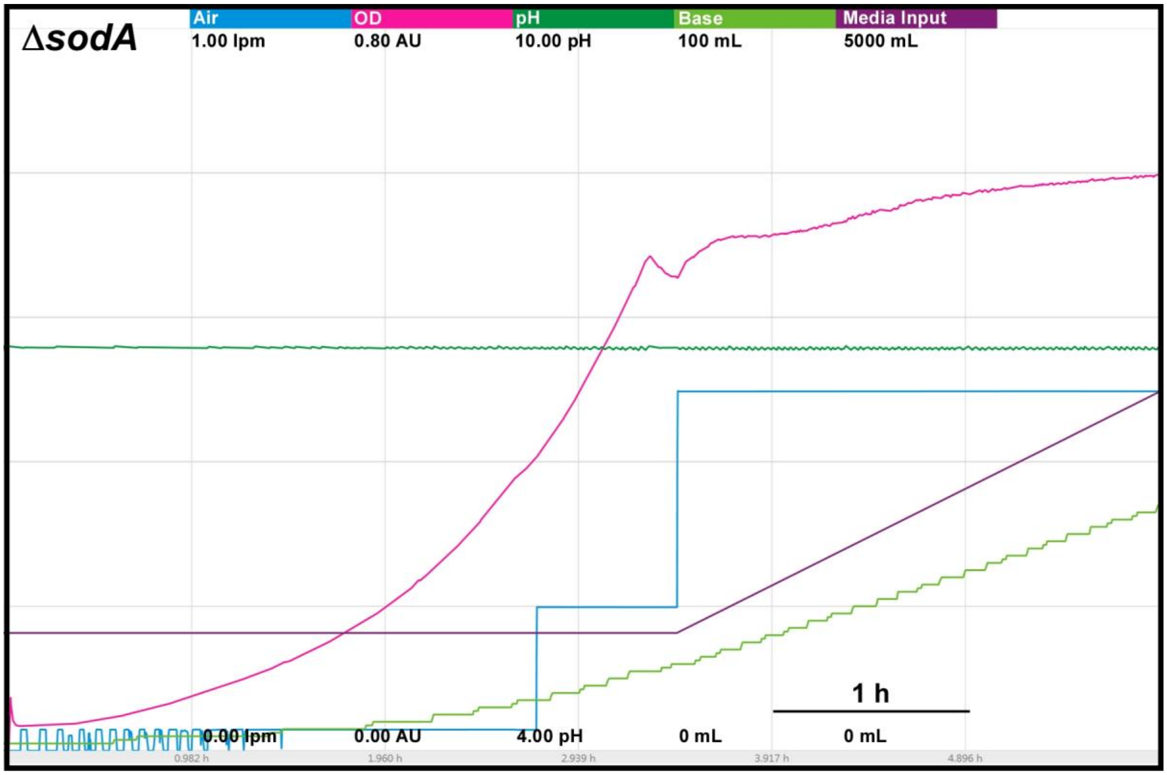


**Figure S5. Aerobic fermentor growth of the Δ*sodA* mutant.**

The Δ*sodA* mutant grown under aerobic fermentor conditions as described previously, without EDTA. Each color represents a different parameter, labeled at the top of the figure. The scale for each parameter is indicated by the values under each respective parameter (minimum at the bottom, maximum at the top). The time scale is indicated by the bar in the bottom right portion of the chart. Representative chart from three replicates.

**A**

| Locus Tag | Annotation | TPM | Log$_2$ Ratio | | |
|---|---|---|---|---|---|
| | | $T_{-20}$ | $T_{10}/T_{-20}$ | $T_{25}/T_{-20}$ | $T_{50}/T_{-20}$ |
| SSA_0363 | *dagA*, sodium:alanine symporter family protein | 120 | 0.23 | **1.26** | **1.22** |
| SSA_0364 | *dctA*, serine/threonine transporter | 102 | 0.30 | **1.16** | **1.14** |
| SSA_0369 | NADP oxidoreductase | 767 | 0.52 | **1.02** | **1.68** |
| SSA_0370 | GNAT family N-acetyltransferase | 1070 | 0.48 | **1.00** | **1.61** |
| SSA_0371 | *gdhA*, NADP-specific glutamate dehydrogenase | 1079 | 0.29 | **0.84** | **1.41** |
| SSA_0385 | *opuAb*, ABC transporter permease/substrate binding protein | 275 | -0.03 | **-1.09** | **-1.74** |
| SSA_0386 | *opuAa*, glycine betaine/L-proline ABC transporter ATP-binding protein | 287 | -0.26 | **-1.32** | **-2.00** |
| SSA_0572 | 2,3-butanediol dehydrogenase | 147 | 0.10 | -0.19 | **-1.75** |
| SSA_0635 | *trpC*, indole-3-glycerol phosphate synthase | 18 | 0.26 | -0.02 | **-1.03** |
| SSA_0637 | *trpB*, tryptophan synthase subunit beta | 55 | **1.22** | **1.25** | **1.38** |
| SSA_0638 | *trpA*, tryptophan synthase subunit alpha | 83 | **1.30** | **1.28** | **1.07** |
| SSA_0703 | citrate synthase | 4 | -0.10 | 0.56 | **1.54** |
| SSA_0704 | NADP-dependent isocitrate dehydrogenase | 7 | -0.51 | 0.44 | **1.03** |
| SSA_0737 | *sagP*, arginine deiminase | 292 | **1.17** | 0.09 | **-1.92** |
| SSA_0738 | *arc*, ornithine carbamoyltransferase | 666 | **0.47** | **-0.93** | **-2.40** |
| SSA_0739 | *arcC*, carbamate kinase | 58 | 0.24 | **-1.45** | **-3.15** |
| SSA_0741 | *arcT*, acetylornithine deacetylase/succinyl-diaminopimelate desuccinylase | 76 | -0.01 | **-1.49** | **-2.99** |
| SSA_0917 | 5-methyltetrahydropteroyltriglutamate-homocysteine methyltransferase | 5 | 0.17 | 0.33 | **1.34** |
| SSA_0921 | *adhB*, alcohol dehydrogenase catalytic domain-containing protein | 128 | -0.04 | **0.75** | **1.23** |
| SSA_1341 | *carB*, carbamoyl-phosphate synthase large subunit | 23 | -0.03 | **0.65** | **1.31** |
| SSA_1342 | *carA*, glutamine-hydrolyzing carbamoyl-phosphate synthase small subunit | 20 | 0.27 | **0.77** | **1.26** |
| SSA_1360 | amino acid ABC transporter ATP-binding protein | 724 | **0.50** | **0.81** | **1.16** |
| SSA_1401 | PLP-dependent aminotransferase family protein | 17 | -0.09 | 0.24 | **1.05** |
| SSA_1439 | *hisK*, hypothetical protein | 73 | **0.62** | **0.78** | **1.41** |
| SSA_1440 | *hisE*, phosphoribosyl-ATP diphosphatase | 75 | 0.16 | 0.47 | **1.11** |
| SSA_1569 | amino acid ABC transporter permease | 112 | **0.77** | **0.91** | **1.01** |
| SSA_1615 | alanine dehydrogenase | 368 | **0.88** | -0.35 | **-3.45** |
| SSA_1621 | amino acid permease | 80 | 0.28 | **-1.19** | **-3.17** |
| SSA_1713 | *serA*, D-3-phosphoglycerate dehydrogenase | 51 | 0.36 | **1.37** | **2.22** |
| SSA_1715 | *serC*, 3-phosphoserine/phosphohydroxythreonine transaminase | 40 | 0.05 | **1.07** | **2.03** |
| SSA_1949 | peptide ABC transporter substrate-binding protein | 182 | **0.40** | **-0.95** | **-2.27** |
| SSA_1950 | peptide ABC transporter ATP-binding protein | 95 | -0.16 | **-0.68** | **-1.38** |
| SSA_1967 | *ilvA*, threonine ammonia-lyase | 58 | 0.21 | **0.81** | **1.32** |
| SSA_1968 | *ilvC*, ketol-acid reductoisomerase | 210 | 0.31 | **0.88** | **1.39** |
| SSA_1969 | *ilvH*, acetolactate synthase small subunit | 108 | -0.09 | 0.57 | **1.09** |
| SSA_1970 | *ilvB*, acetolactate synthase large subunit | 114 | -0.34 | **0.49** | **1.01** |
| SSA_2039 | amino acid permease | 23 | 0.01 | **-0.97** | **-1.18** |
| SSA_2141 | *argH*, argininosuccinate lyase | 14 | 0.06 | 0.39 | **1.24** |

| Log$_2$ Ratio |
|---|
| 2.5 |
| 2.0 |
| 1.5 |
| 1.0 |
| 0.5 |
| 0.0 |
| -0.5 |
| -1.0 |
| -1.5 |
| -2.0 |
| -2.5 |
| -3.0 |
| -3.5 |
| -4.0 |
| -4.5 |

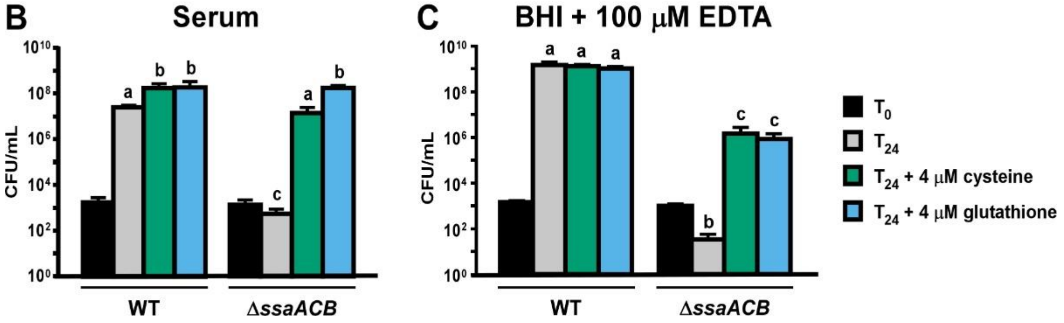


**Figure S6. Impact of Mn depletion on amino acid transport and synthesis**

Expression of amino acid transport and synthesis genes in the Δ*ssaACB* mutant are depicted with their average TPM at $T_{-20}$ and $\log_2$ fold change values for each post-EDTA time point. Only genes with $\log_2$ fold change values ≥ |1| are depicted in this chart. For all genes, see Table S1. TPM values greater than 1000 are full saturation (green). Positive $\log_2$ fold change values (red) are upregulated in post-EDTA samples as compared to $T_{-20}$, while negative values (blue) are downregulated. Values in bold are significant by adjusted *P*-value (≤ 0.05). WT and Δ*ssaACB* cells were grown for 24 hours at 12% $O_2$ in either (**B**) pooled rabbit serum or (**C**) BHI + 100 µM EDTA with 4 µM of either cysteine or glutathione added. The means and standard deviations of at least three replicates are displayed. Significance was determined by repeated measures ANOVA with a Tukey-Kramer multiple comparisons test. $T_{24}$ bars with the same letter are not significantly different from each other. $T_0$ values from each experiment were compared to each other by Student's two tailed t-test and found to be not significantly different.

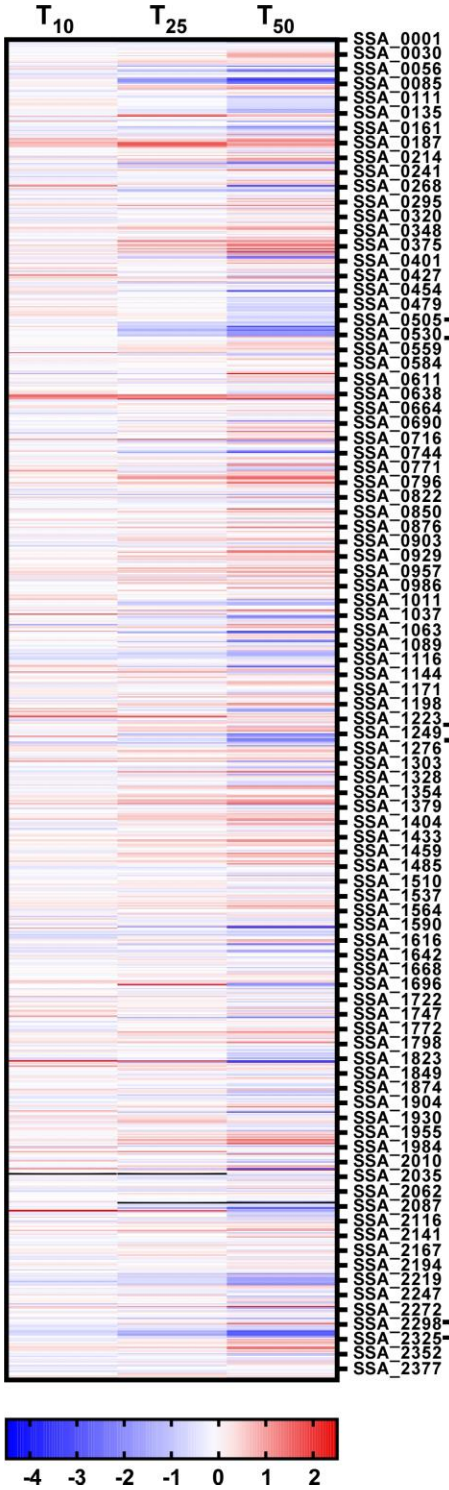


| Locus Tag | Annotation | TPM | Log₂ Ratio | | |
|---|---|---|---|---|---|
| | | $T_{-20}$ | $T_{10}/T_{-20}$ | $T_{25}/T_{-20}$ | $T_{50}/T_{-20}$ |
| **Ethanolamine Utilization Proteins** | | | | | |
| SSA_0507 | hypothetical protein | 539 | -0.36 | **-0.90** | **-1.06** |
| SSA_0508 | Phosphoserine phosphatase | 63 | **0.67** | **-0.87** | **-2.07** |
| SSA_0509 | Propanediol utilization protein, PduV/EutP | 84 | 0.17 | **-1.15** | **-2.28** |
| SSA_0510 | L-threonine-O-3-phosphate decarboxylase, CobD | 157 | 0.12 | **-0.61** | **-0.65** |
| SSA_0511 | L-threonine kinase, PduX | 145 | -0.04 | **-0.66** | **-0.61** |
| SSA_0512 | Phosphoribosyltransferase, CobT | 169 | 0.01 | **-0.69** | **-0.62** |
| SSA_0513 | Ethanolamine utilization, CobO | 165 | -0.03 | **-0.80** | **-0.52** |
| SSA_0514 | Ethanol dehydrogenase, EutG | 160 | **0.47** | **-0.97** | **-3.52** |
| SSA_0515 | Ethanolamine utilization protein, EutS | 1683 | 0.07 | **-0.83** | **-2.06** |
| SSA_0516 | Ethanolamine two-component response regulator, EutV | 1069 | 0.07 | **-0.79** | **-1.84** |
| SSA_0517 | Ethanolamine transduction histidine kinase, EutW | 957 | 0.27 | **-0.80** | **-1.84** |
| SSA_0518 | Ethanolamine utilization protein EutA | 48 | **0.71** | **-0.97** | **-1.82** |
| SSA_0519 | Ethanolamine ammonia-lyase large subunit, EutB | 62 | 0.13 | **-1.26** | **-1.97** |
| SSA_0520 | Ethanolamine ammonia-lyase small subunit, EutC | 50 | 0.10 | **-1.02** | **-1.72** |
| SSA_0521 | Ethanolamine utilization, EutL | 85 | -0.24 | **-1.42** | **-2.55** |
| SSA_0522 | Ethanolamine utilization, EutK | 52 | 0.24 | **-0.77** | **-1.57** |
| SSA_0523 | Acetaldehyde dehydrogenase,EutE | 70 | 0.02 | **-0.88** | **-1.66** |
| SSA_0524 | Ethanolamine utilization protein, PduA/PduJ | 79 | -0.15 | **-0.91** | **-1.58** |
| SSA_0525 | Ethanolamine utilization protein, PduA/PduJ | 115 | -0.10 | **-0.77** | **-1.88** |
| SSA_0526 | Ethanolamine utilization, EutT | 95 | -0.13 | **-0.92** | **-1.87** |
| SSA_0527 | Ethanolamine utilization, EutD | 99 | -0.06 | **-0.84** | **-1.86** |
| SSA_0528 | hypothetical protein | 79 | -0.03 | **-0.83** | **-1.79** |
| SSA_0529 | Ethanolamine utilization, EutN | 96 | -0.14 | -0.72 | **-1.76** |
| SSA_0530 | Ethanolamine permease, EutH | 122 | -0.21 | **-1.10** | **-1.94** |
| SSA_0531 | Ethanolamine utilization protein, EutQ | 161 | -0.35 | **-1.12** | **-2.26** |
| **CRISPR-Associated Proteins** | | | | | |
| SSA_1246 | CRISPR-associated protein Csm6 | 130 | 0.02 | **-1.31** | **-2.88** |
| SSA_1247 | CRISPR-associated protein, Csm5 family | 74 | -0.33 | **-1.33** | **-2.38** |
| SSA_1248 | CRISPR-associated RAMP protein, Csm4 family | 67 | -0.27 | **-1.21** | **-2.47** |
| SSA_1249 | CRISPR-associated RAMP Csm3 | 61 | -0.08 | **-1.20** | **-2.53** |
| SSA_1250 | CRISPR-associated protein, Csm2 family | 54 | -0.06 | **-1.01** | **-2.03** |
| SSA_1251 | CRISPR-associated protein, Csm1 family | 42 | 0.22 | -0.61 | **-2.01** |
| SSA_1252 | CRISPR-associated endoribonuclease Cas6 | 101 | **1.45** | 0.35 | **-1.88** |
| SSA_1253 | CRISPR-associated protein Cas2 | 8 | *-0.46* | -0.38 | -1.15 |
| SSA_1254 | hypothetical protein | 3 | *-0.28* | *-0.29* | -1.06 |
| SSA_1255 | CRISPR-associated protein Cas1 | 5 | *-0.49* | -0.94 | **-1.68** |
| **Type IV Pilus Proteins** | | | | | |
| SSA_2302 | Prepilin peptidase | 396 | **-0.36** | **-0.45** | **-0.70** |
| SSA_2303 | hypothetical protein | 499 | **-0.33** | **-0.47** | **-0.88** |
| SSA_2304 | hypothetical protein | 645 | **-0.37** | **-0.55** | **-0.68** |
| SSA_2305 | AAA ATPase | 440 | **-0.40** | **-0.70** | **-1.20** |
| SSA_2307 | Type IV pilus biogenesis protein PilO | 48 | **-0.69** | **-1.15** | **-2.83** |
| SSA_2308 | Type IV pilus biogenesis protein PilN | 28 | -0.62 | **-0.89** | **-1.92** |
| SSA_2309 | Type IV pilus biogenesis protein PilM | 44 | **-0.62** | **-1.45** | **-2.74** |
| SSA_2310 | hypothetical protein | 62 | **-0.87** | **-1.46** | **-3.10** |
| SSA_2311 | hypothetical protein | 56 | **-0.62** | **-1.38** | **-2.75** |
| SSA_2312 | hypothetical protein | 35 | -0.72 | **-1.05** | **-2.62** |
| SSA_2313 | Type IV pilin PilA | 183 | -0.48 | **-1.61** | **-2.82** |
| SSA_2314 | Type IV pilin PilA | 311 | **-0.61** | **-1.66** | **-2.96** |
| SSA_2315 | Type IV pilus biogenesis protein PilE | 991 | -0.35 | **-1.23** | **-2.42** |
| SSA_2316 | Type IV fimbrial assembly protein PilC | 107 | -0.49 | **-1.42** | **-2.55** |
| SSA_2317 | Twitching motility protein PilT | 107 | **-0.91** | **-1.36** | **-2.60** |
| SSA_2318 | Type IV fimbrial assembly, ATPase PilB | 406 | **-0.78** | **-1.63** | **-3.21** |

**Figure S7. Transcriptomic heatmap of Δ*ssaACB* aerobic fermentor grown cells**

Heatmap displaying the $\log_2$ fold change values of each gene at the time indicated as compared to $T_{-20}$. Positive $\log_2$ fold change values (red) are upregulated in later samples as compared to $T_{-20}$, while negative values (blue) are downregulated. Select genes are depicted with their average TPM at $T_{-20}$ and $\log_2$ fold change values for each post-EDTA time point. TPM values greater than 1000 are full saturation (green). $\text{Log}_2$ fold change values follow the same color scale as depicted in the heatmap. Values in bold are significant by adjusted $P$-value ($\leq 0.05$).

#### Supplementary Methods

##### Fermentor set up

BHI was prepared in a polypropylene carboy to a final volume of 5 L. Antifoam (Sigma) was added to 25 ppm and the carboy was autoclaved for 2 h at 128°C. Aeration was achieved by the controlled addition of compressed $N_2$ and/or air delivered through a ring sparger, augmented by stirring at 250 rpm; vessel temperature was controlled via an external heating blanket. The dissolved oxygen (DO), OD, and temperature were also measured by indwelling probes. Constant volume in the vessel was maintained by the placement of the opening of a harvest tube at the 800 mL level.

##### Quantitative reverse transcriptase polymerase chain reaction

RNA was collected as described in the main text. cDNA libraries were created using SensiFAST cDNA Synthesis Kit (Bioline). Control reactions without reverse transcriptase were conducted to confirm the absence of contaminating DNA in all samples. qRT-PCR was performed using SYBR Green Supermix (Applied Biosystems) on an Applied Biosystems 7500 Fast Real Time PCR System using the primers listed in Table S3. Relative gene expression was analyzed using the $2^{-\Delta\Delta CT}$ method (Livak and Schmittgen, 2001) with *gapA* serving as the internal control (Rodriguez et al., 2011).

##### Heatmap generation

The heatmap was generated in GraphPad Prism v. 8.2.0 (graphpad.com) using the $\log_2$ fold change values of the RNA-Seq data calculated using DESeq2 in Geneious, as described in the main text.

##### Putative *cre* site identification

Putative *cre* sites identified in the SK36 genome by RegPrecise (https://enigma.lbl.gov/regprecise/; RRID:SCR_002149) (Novichkov et al., 2013) and listed within the "propagated regulon" collection (http://regprecise.sbpdiscovery.org:8080/WebRegPrecise/regulon.jsp?regulon_id=35148) were collected. Further analyses were performed by obtaining non-overlapping sequences ≤250 bp in length upstream of all SK36 genes using RSAT (http://rsat.sb-roscoff.fr/; RRID:SCR_008560) (van Helden et al., 2000; Nguyen et al., 2018), then searching for putative *Streptococcus suis* pseudopalindromic *cre* and *cre*2 motifs (Willenborg et al., 2014)in these sequences using FIMO from MEME Suite (http://meme-suite.org/doc/fimo.html; RRID:SCR_001783) (Grant et al., 2011). Motifs used for each search, as well as scores given for the RegPrecise and FIMO outputs, are listed in Table S2. Due to the variable length of the $cre_{var}$ sites (Yang et al., 2017), seqinR v 3.6-1 (Charif and Lobry, 2007) was used for this search. The FIMO cutoff was set to $P$-value $\leq 10^{-4}$. Putative sites located within 10 bp of the start site of the corresponding gene were removed from the list.

**Supplementary References**

Charif, D., and Lobry, J.R. (2007). "SeqinR 1.0-2: A Contributed Package to the R Project for Statistical Computing Devoted to Biological Sequences Retrieval and Analysis," in *Structural Approaches to Sequence Evolution: Molecules, Networks, Populations,* eds. U. Bastolla, M. Porto, H.E. Roman & M. Vendruscolo. (Berlin, Heidelberg: Springer Berlin Heidelberg), 207-232.

Grant, C.E., Bailey, T.L., and Noble, W.S. (2011). FIMO: scanning for occurrences of a given motif. *Bioinformatics* 27(7)**,** 1017-1018. doi: 10.1093/bioinformatics/btr064.

Livak, K.J., and Schmittgen, T.D. (2001). Analysis of relative gene expression data using real-time quantitative PCR and the $2^{(-\Delta\Delta CT)}$ method. *Methods* 25(4)**,** 402-408. doi: 10.1006/meth.2001.1262.

Nguyen, N.T.T., Contreras-Moreira, B., Castro-Mondragon, J.A., Santana-Garcia, W., Ossio, R., Robles-Espinoza, C.D., et al. (2018). RSAT 2018: regulatory sequence analysis tools 20th anniversary. *Nucleic Acids Res* 46(W1)**,** W209-w214. doi: 10.1093/nar/gky317.

Novichkov, P.S., Kazakov, A.E., Ravcheev, D.A., Leyn, S.A., Kovaleva, G.Y., Sutormin, R.A., et al. (2013). RegPrecise 3.0 - A resource for genome-scale exploration of transcriptional regulation in bacteria. *BMC Genomics* 14**,** 745. doi: 10.1186/1471-2164-14-745.

Rodriguez, A.M., Callahan, J.E., Fawcett, P., Ge, X., Xu, P., and Kitten, T. (2011). Physiological and molecular characterization of genetic competence in *Streptococcus sanguinis*. *Mol Oral Microbiol* 26(2)**,** 99-116. doi: 10.1111/j.2041-1014.2011.00606.x.

van Helden, J., André, B., and Collado-Vides, J. (2000). A web site for the computational analysis of yeast regulatory sequences. *Yeast* 16(2)**,** 177-187. doi: 10.1002/(sici)1097-0061(20000130)16:2.

Willenborg, J., de Greeff, A., Jarek, M., Valentin-Weigand, P., and Goethe, R. (2014). The CcpA regulon of *Streptococcus suis* reveals novel insights into the regulation of the streptococcal central carbon metabolism by binding of CcpA to two distinct binding motifs. *Mol Microbiol* 92(1)**,** 61-83. doi: 10.1111/mmi.12537.

Yang, Y., Zhang, L., Huang, H., Yang, C., Yang, S., Gu, Y., et al. (2017). A flexible binding site architecture provides new insights into CcpA global regulation in gram-positive bacteria. *mBio* 8(1). doi: 10.1128/mBio.02004-16.
